# DOT1L interaction partner AF10 controls patterning of H3K79 methylation and RNA polymerase II to maintain cell identity

**DOI:** 10.1101/2020.12.17.423347

**Authors:** Coral K. Wille, Edwin N. Neumann, Aniruddha J. Deshpande, Rupa Sridharan

**Author notes:** Corresponding author: Dr. Rupa Sridharan, Wisconsin Institute for Discovery, University of Wisconsin, 330 North Orchard Street, Room 2118, Madison, WI 53715.

## Abstract

Histone 3 Lysine 79 methylation (H3K79me) is enriched on gene bodies proportional to gene expression levels, and serves as a strong barrier for reprogramming of somatic cells to induced pluripotent stem cells (iPSCs). DOT1L is the sole histone methyltransferase that deposits all three orders - mono, di, and tri methylation – at H3K79. Here we leverage genetic and chemical approaches to parse the specific functions of the higher orders of H3K79me in maintaining cell identity. DOT1L interacts with AF10 (*Mllt10*) which recognizes unmodified H3K27 and boosts higher order H3K79 methylation. AF10 deletion evicts higher order H3K79me2/3, and reorganizes H3K79me1 to the transcription start site, to facilitate iPSC formation in the absence of steady state transcriptional changes. Instead, AF10 loss redistributes RNA polymerase II to a uniquely pluripotent pattern at highly expressed, rapidly transcribed housekeeping genes. Taken together, we reveal a mechanism for higher order histone methylation located at the gene body in reinforcing cell identity.

## INTRODUCTION

Histone modifications shape the epigenome to be responsive to the transcriptional machinery that controls gene expression. While the function of histone modifications that determine chromatin compaction to allow transcription factor binding is intuitive, those that accompany transcriptional elongation have been obscure. Histone acetylation on most residues de-compacts chromatin to facilitate transcription, but the effects of histone methylation vary depending on the particular residue. Further methylation can occur to three orders – me1, me2 and me3, with potentially different effects on transcriptional output. The enzymes that write or erase histone modifications are found in complexes with proteins that can regulate their activity to higher order methylation.

DOT1L is the exclusive methyltransferase of all degrees of histone 3 lysine (K) 79 methylation (H3K79me1, me2, and me3), a modification found on gene bodies [1]. DOT1L is a distributive enzyme and longer residence time leads to greater degree of methylation [2]. Thus, the same loci tend to be enriched for H3K79me1, me2 and me3. While the individual functions of each modification remain unknown, all levels of H3K79me are enriched after transcription has begun [3]. H3K79me2 in particular has been correlated with a rapid transcriptional rate [4–6].

DOT1L has a DNA-binding domain and an H2BK120 ubiquitin interaction domain that stabilizes DOT1L-histone binding and promotes methylation of H3K79 [7,8]. DOT1L also forms complexes with numerous associated proteins that have histone interaction domains and promote higher order methylation. Among these, AF10 and AF17 contain a PZP module that interacts with unmodified H3K27 [9]. AF10 functions as a co-factor to increase DOT1L catalytic activity for both H3K79me2 and me3 [10]. AF9 and ENL contain a YEATS domain that binds to H3K9 and H3K27 acylations [11,12], and their depletion decreases H3K79me3 without an effect on H3K79me2 [13]. Thus, DOT1L complex composition could greatly affect both the genomic loci to which DOT1L is targeted and the degree of H3K79 methylation that is output.

H3K79me2 is the most differentially enriched histone modification in mouse embryonic fibroblasts (MEFs) compared to embryonic stem cells (ESCs), with a 7 fold greater abundance in MEFs [14]. ESCs uniquely shuttle DOT1L to the cytoplasm [15]. Inhibition or loss of DOT1L greatly enhances the reprogramming of induced pluripotent stem cells from somatic cells [16–18]. Thus low DOT1L activity is coincident with, and even promotes, a gain of potency [19].

Here we define the role of the degrees of H3K79 methylation in the non-transformed and highly dynamic environment of somatic cell reprogramming to induced pluripotent stem cells (iPSCs). We find that DOT1L maintains cellular identity through AF10, but not AF9 or ENL. Deletion of AF10 results in removal of H3K79me2 and H3K79me3, and increases reprogramming efficiency to almost half the levels seen by DOT1L catalytic inhibition where all H3K79me is lost. Structure-function analysis of AF10 reveals that the AF10-histone interaction is dispensable, but the AF10-DOT1L interaction is required for maintenance of somatic cell fate. Although H3K79me1/2 is enriched at thousands of genes, AF10 depletion results in few transcriptional changes, the majority of which are not congruent with expression in ESCs.

Instead, deletion of AF10 results in an increase of H3K79me1 with a concomitant reduction in RNAPII specifically at the transcription start site. Strikingly, high H3K79me1-lower RNAPII is the pattern found in ESCs. Therefore, AF10 functions as a DOT1L enzymatic cofactor in maintaining RNAPII patterning in preserving cell identity.

## RESULTS

### Higher order H3K79 methylation affects reprogramming efficiency

To interrogate the role of H3K79 methylation in cell identity, we used the system of reprogramming somatic cells to induced pluripotent stem cells where DOT1L is a strong barrier for cell fate change [16–18]. We examined the expression of DOT1L interacting partners AF10, AF9 and ENL in mouse somatic cells (MEFs), cells on day 2 and day 4 of reprogramming, and ESCs (Fig 1A). All assessed DOT1L associated factors were expressed throughout reprogramming, yet with different patterns from both bulk (Fig 1A) and single cell RNA-seq (scRNA-seq) [20] (Fig S1A). DOT1L itself was expressed at low levels in few cells regardless of pluripotency status (Fig S1A), while being largely co-expressed with ENL in MEFs, AF9 during reprogramming, and AF10 in ESCs (Fig S1B).

**Figure 1.**
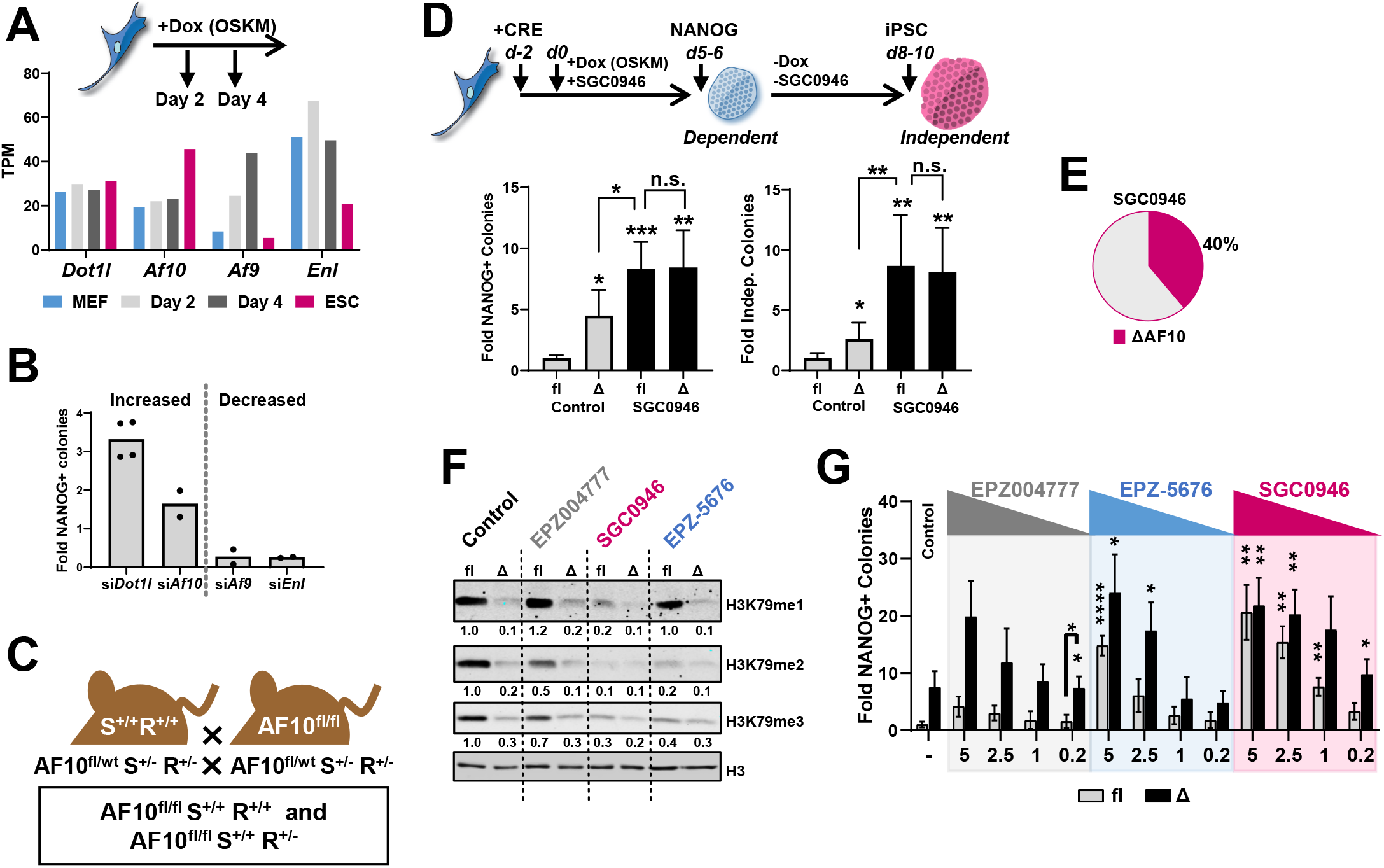
Higher order H3K79 methylation affects reprogramming efficiency. A. Top: MEFs, cells on days 2 and 4 of reprogramming, and ESCs were transcriptionally profiled. OSKM = Oct4, Sox2, Klf4, and c-Myc induced by doxycycline (dox) treatment. Bottom: Expression of DOT1L associated factors determined by bulk RNA-Seq [18] measured as transcripts per million (TPM). B. Reprogramming upon siRNA depletion of indicated genes. Colonies obtained in non-targeting control set to 1. Data are the mean of at least 2 biological replicates, depicted as dots. C. Breeding scheme to generate conditional AF10 mice with inducible reprogramming factors. AF10fl/flS+/+R+/+ and AF10fl/flS+/+R+/- were used for all studies. fl = flox, wt = wild-type allele, S = Stemcca cassette (dox inducible OSKM), R = reverse tetracycline-controlled transactivator. D. Top: Reprogramming scheme – mouse embryonic fibroblasts (MEFs) were infected with empty adenovirus (fl) or adenovirus-CRE to delete AF10 (ΔAF10). Reprogramming was initiated 2 days (d) later with dox, in the presence and absence of DOT1Li treatment. OSKM transgene dependent reprogramming efficiency was measured by the number of NANOG+ after 5-6 days of continuous dox and drug treatment. Transgene independent colonies sustained NANOG 2-4 days after dox and drug withdrawal. Bottom: OSKM transgene dependent (Left) and independent (Right) reprogramming. Colonies obtained in fl treated with control set to 1. Data are the mean + S.D. (n = 3). ***P<0.001, **P<0.01, *P<0.05, and not significant (n.s.) P>0.05 by Welch’s ANOVA with post-hoc pairwise t-tests using the Holm correction (significance against fl control displayed unless indicated). E. Average efficiency of ΔAF10 in transgene dependent reprogramming. Colonies obtained in ΔAF10+DOT1Li set to 100% (n=8). F. Immunoblot of H3K79me1, me2, and me3 in AF10 fl or Cre deleted (Δ) conditions treated with 5 μM of DOT1L inhibitors on day 3 of reprogramming. Numbers (below) indicate signal relative to total H3 (fl treated with Control set to1). G. NANOG+ colonies on day 5-7 of reprogramming of control fl (gray bars) and AF10 deleted Δ (black bars) cells treated with decreasing amounts (5, 2.5, 1, 0.2 μM) of DOT1L inhibitors: EPZ004777 (gray), SGC0946 (red), and EPZ-5676 (blue). Colonies obtained in fl cells control treated set to 1. Data are the mean + S.D. (n = 3). ****P<0.0001, **P<0.01, *P<0.05, and not significant (n.s.) P>0.05 by Welch’s ANOVA with post-hoc pairwise t-tests using the Holm correction (only significant conditions against fl with control treatment, or otherwise labeled, are denoted).

Given this variation throughout reprogramming, we tested the effect of DOT1L associated factors during the process. We initiated reprogramming in MEFs isolated from a reprogrammable mouse model in which the Yamanaka factors Oct4, Sox2, c-Myc and Klf4 are expressed from a single cassette under the control of a doxycycline inducible promoter. Each DOT1L associated factor was depleted using siRNA (Fig S1C) concurrent with the initiation of reprogramming. Successful iPSC generation was measured by the expression of the pluripotency protein NANOG. While depletion of AF9 and ENL both decreased reprogramming efficiency, like DOT1L, AF10 was a barrier for reprogramming (Fig 1B).

To better elucidate the role of AF10 in reprogramming, AF10 conditional mice [10] were bred with the inducible reprogrammable mouse line (Fig 1C). In somatic cells isolated from the resulting AF10 reprogrammable mouse line, both the transition to pluripotency (Fig 1D) and AF10 function (Fig S1D) can be temporally controlled. We first compared the reprogramming efficiency of AF10 deletion (Δ) to the undeleted (fl) control. ΔAF10 increased NANOG+ colonies by greater than 4-fold in the presence of exogenous reprogramming factor expression (Fig 1D, Left). Deletion of AF10 also increased the formation of *bona fide* colonies that remained after the withdrawal of doxycycline-driven reprogramming factors, a more stringent test of pluripotency acquisition (Fig 1D, Right). Further, iPSCs that were picked and expanded from control treated (fl) and CRE-deleted (Δ)AF10 reprogramming cells maintained pluripotent colony morphology and expressed similar levels of the pluripotency the factor OCT4 (Fig S1E) after multiple passages without dox.

Since both AF10 and DOT1L can have pleiotropic functions, we next asked if AF10 functioned in the same pathway as DOT1L in pluripotency acquisition. We used a potent DOT1L catalytic inhibitor SGC0946 [21] in combination with CRE-recombinase mediated deletion of AF10. Approximately 2-fold more colonies were obtained in SGC0946 as compared to ΔAF10 alone (Fig 1D, black bars). A near complete loss in AF10 (Fig S1D) contributes about 40% of the DOT1Li phenotype in iPSC generation (Figs 1E and S1F). Thus, deletion of AF10 functions in the same pathway to DOT1L inhibition during reprogramming.

We further tested the relationship between degrees of H3K79 methylation and reprogramming efficiency using other commercially available inhibitors of DOT1L that have different efficacy: EPZ0044777, which is structurally similar to SGC0946 but 10-fold less potent, and EPZ-5676, a drug with similar potency to SGC0946 but with increased drug properties for use in clinic [21–24]. We first tested whether this reported difference in potency between the DOT1L inhibitors was also observed in the reprogramming system. On day 3 of reprogramming, in control MEFs (fl-AF10 flox), EPZ004777 failed to remove H3K79me1 and marginally depleted H3K79me2 and me3, while EPZ-5676 retained only H3K79me1 compared to the most potent inhibitor SGC0946 (Fig 1F). Consequently, EPZ004777 treated fl control had 20% of the reprogrammed colonies, while EPZ-5676 treated fl control achieved about 70% of the reprogramming efficiency of SGC0946 (Fig 1G). Upon AF10 deletion, the number of iPSC colonies obtained with EPZ004777 and EPZ-5676 was equal to that seen with SGC0946 (Fig 1G) when H3K79me1/2/3 removal was comparable (Fig 1F). Further, there was a dose-dependent response in AF10 deletion effects-for example, at the highest concentration (5 μM) of SGC0946, there was only a 1.1-fold increase of iPSC colonies as compared to the lower concentration (0.2 μM) which showed a 3.0-fold increase (Fig 1G). This boost in reprogramming efficiency upon AF10 deletion largely corresponded to removal of residual H3K79 methylation.

To identify if AF10 has a temporal effect during pluripotency acquisition, AF10 was deleted coincident with reprogramming factor induction (day 0), at the midpoint (day 2) and later (day 4) in a reprogramming timecourse, which assessed NANOG+ colony formation on day 6 (Fig S1G). Similar to DOT1L inhibition [17,18,25], loss of AF10 most potently increased early and mid-reprogramming (Fig S1G). Taken together, AF10 collaborates with DOT1L to maintain somatic identity and its loss facilitates cell identity conversion.

### AF10 deletion has few transcriptional effects

To determine how AF10 deletion enabled acquisition of pluripotency, we performed RNA-seq experiments. Based on the timecourse of maximum iPSC generation (Fig S1G), we performed RNA-seq on day 4 of reprogramming of cells that were treated with control or CRE recombinase at the initiation of reprogramming factors, compared to SGC0946, henceforth called DOT1Li. The ΔAF10 expression profile is most correlated with DOT1Li treated cells, and not ESCs as expected for a sample from the midpoint of the process (Fig 2A-B).

**Figure 2.**
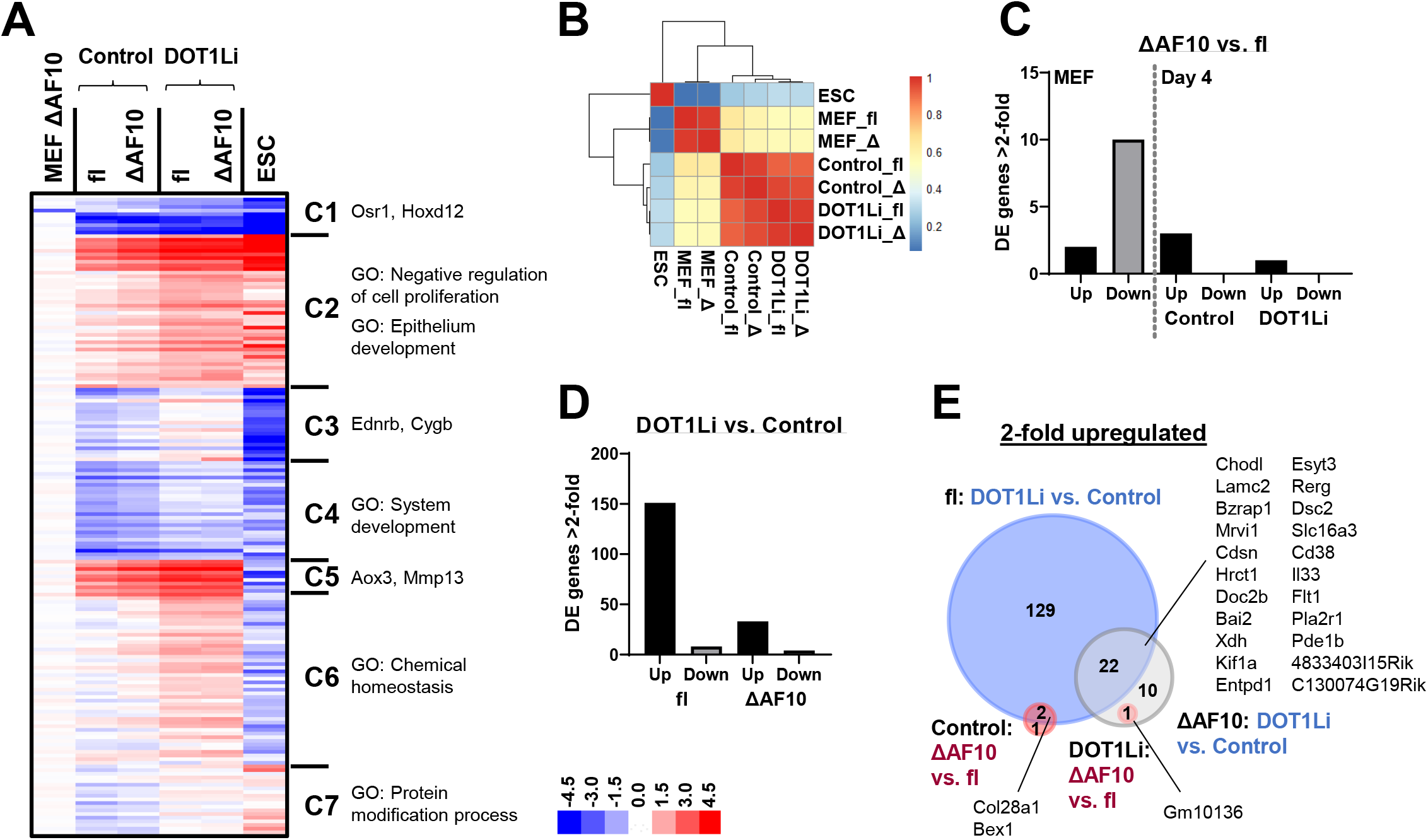
AF10 deletion has few transcriptional effects. A. k-means clustered heatmap of Log2 fold change in transcripts per million (TPM) relative to fl MEFs of differentially expressed (DE) genes from all conditions. B. Heatmap of Pearson correlations with hierarchical clustering of TPM values of DE genes from all conditions. C-D. Number of DE genes (2-fold or more with a posterior probability of differential expression 0.95 or greater) upregulated (black) or downregulated (gray) in: C. ΔAF10 vs. fl cells: in MEFs (left) or day 4 of reprogramming (right, as indicated); D. DOT1Li vs. Control on day 4 of reprogramming of: fl cells (left) and ΔAF10 cells (right). E. Overlap of upregulated genes on day 4 of reprogramming in: DOT1Li vs. Control in fl (blue), DOT1Li vs. Control in ΔAF10 (gray), ΔAF10 vs. fl in control (red), and ΔAF10 vs. fl in DOT1Li (pink) conditions.

To observe the trend in functional categories that changed dynamically, we generated a heatmap of differentially expressed (DE) genes from all conditions. Of the two clusters (C1 and C2) that were altered by DOT1Li to resemble ESCs (Fig 2A), C2 contained genes that functioned in negative regulation of cellular proliferation and epithelium development (Fig S2A). However, most of the transcriptional changes are not representative of ESC gene expression (Fig 2A). In reprogramming unique clusters C3-C6, the magnitude of gene expression changes in ΔAF10 is lower than that in DOT1Li. Cluster 7 contains protein modification process genes that are modestly upregulated upon DOT1Li and in ESCs, but are downregulated in AF10 deleted cells (Fig S2B). None of the clusters showed a major decrease in MEF specific genes in contrast to observations in AF10 depleted human reprogramming conditions [26]. Thus, suppression of somatic expression by AF10 deletion is not the primary mode of facilitating iPSC generation.

Although deletion of AF10 positions cells closer to DOT1Li treatment (Fig 2B), few genes are significantly DE measured as 2-fold or more fold-change in expression, with a posterior probability of being DE greater than 0.95 (Methods). At the starting point in MEFs, fewer than 15 genes were DE (Fig 2C). On day 4 of reprogramming in control treated cells, ΔAF10 resulted in the upregulation of 3 genes relative to control, and there was a single upregulated gene in ΔAF10 in cells treated with DOT1Li (Fig 2C) further demonstrating that deletion of AF10 does not add to DOT1L inhibition. Treatment of cells with DOT1Li had the greatest effect with many more genes upregulated (153) than downregulated (8) relative to control treatment (Fig 2D). DOT1Li treatment of ΔAF10 cells produced 33 upregulated genes and 4 downregulated (Fig 2D). Twenty-two genes were upregulated by DOT1Li in both fl and ΔAF10 cells (Fig 2E), and only *Hoxd12* and *Osr1* were commonly downregulated. Of note, we have demonstrated that reduction of *Hoxd12* did not contribute to the DOT1Li reprogramming phenotype [18]. Overall, the loss of AF10 yielded almost no significant transcriptional changes at the steady state mRNA level although it lessened the DOT1Li transcriptional effect (Fig 2D). Therefore, we examined histone modification circuitry effects that may not yield immediate changes in gene expression by which ΔAF10 could contribute to pluripotency acquisition.

### AF10 enforces cellular identity through DOT1L interaction

AF10 is comprised of an N-terminal PHD finger-Zn knuckle-PHD finger (PZP) module that interacts with unmodified histone H3K27 [9] and a C-terminal octapeptide motif–leucine-zipper (OMLZ) domain that interacts with DOT1L (Fig 3A). To uncover which of these domains may be important for cellular identity we performed structure function analysis.

**Figure 3.**
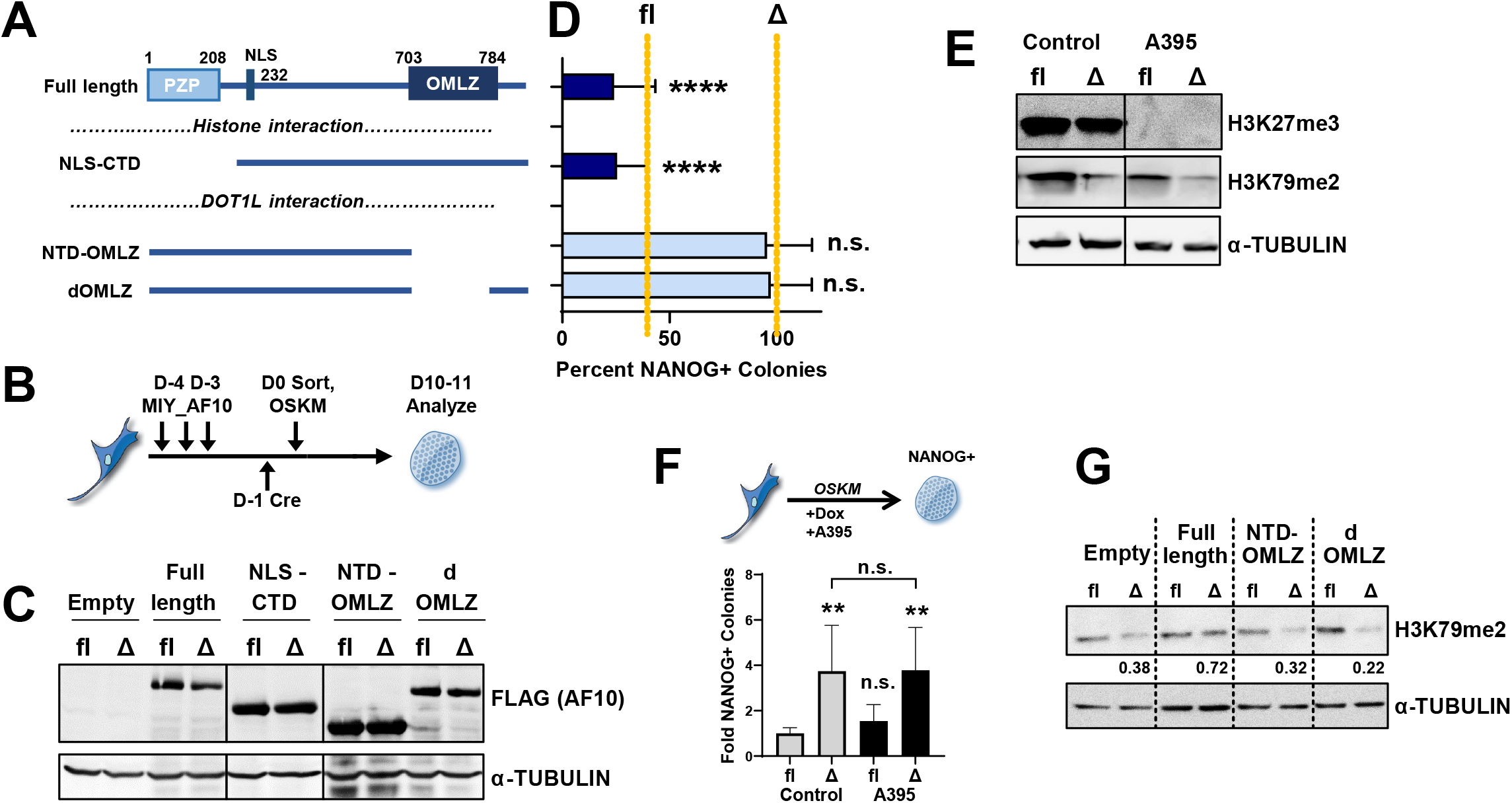
AF10 enforces cellular identity through DOT1L interaction. A. AF10 contains two major functional domains: 1) PHD finger-Zn knuckle-PHD finger (PZP) module and 2) octapeptide motif-leucine zipper (OMLZ). Domain mutants that ablate histone interaction (NLS-CTD) or DOT1L interaction (NTD-OMLZ and dOMLZ) are depicted. B. MEFs were infected 3 times with retroviruses containing AF10 domain mutants over 3-4 days prior to reprogramming. Endogenous AF10 was deleted with Cre recombinase or cells treated with control adenovirus on day-1. The following day, cells were sorted by flow cytometry, plated, and reprogramming (OSKM) was initiated. C. Immunoblot of AF10 domain mutant expression on day 5 of reprogramming in control (fl) and AF10 deleted (Δ) cells. D. Reprogramming efficiency of AF10-Cre deleted cells (ΔAF10), expressing the indicated exogenous AF10 domain mutants (Fig 4A). Reprogramming of wild type cells (fl) and ΔAF10 infected with empty vector control indicated by dotted lines. Colonies in ΔAF10 infected with empty vector control set to 100%. Data are mean + S.D. (n = 4). ****P<0.0001 and not significant (n.s.) P>0.05 by one-way ANOVA with Tukey’s multiple comparison test post-hoc. Significance against ΔAF10 with empty vector control (dotted line) displayed. E. Immunoblot of H3K79me2, H3K27me3, and α-TUBULIN in cells treated with control or A395 (EED inhibitor) on day 5 of reprogramming. F. Transgene dependent reprogramming of fl or ΔAF10 cells, treated with A395 or control. Data are the mean + S.D. (n = 5). **P<0.01 and not significant (n.s.) P>0.05 by Welch’s ANOVA with post-hoc pairwise t-tests using the Holm correction (significance against fl control displayed unless indicated). G. Immunoblot of H3K79me2 in cells with endogenous (fl) or deleted (Δ) AF10, exogenously expressing AF10 domain mutants, on day 5 of reprogramming. H3K79me2 signal normalized to α-TUBULIN was calculated in ΔAF10 relative to matched fl conditions (Bottom).

We generated one mutant of AF10 that cannot interact with histones (NLS-CTD) and two mutants that omitted either the OMLZ domain alone (delta-dOMLZ) or in combination with the C-terminal domain (NTD-OMLZ) to prevent interaction with DOT1L (Fig 3A) in a vector with a fluorescent reporter. AF10 mutants were individually transduced into AF10 conditional reprogrammable MEFs followed by deletion of endogenous AF10 or control treatment (Fig 3B). MEFs containing exogenous AF10 mutants were isolated with flow cytometry (Fig S3A) and reprogramming was initiated (Fig 3B). Equivalent levels of each mutant protein were observed in the sorted reprogramming cells (Fig 3C). Therefore, each reprogramming condition expressed either exogenous AF10 mutants alone (Δ) (Fig 3D, blue bars) or mutant proteins in addition to the endogenous AF10 (fl) (Fig S3B, blue bars). While deletion of AF10 enhanced reprogramming (Figs 3D and S3B dotted line), expression of exogenous full length AF10 reduced the number of NANOG+ colonies during reprogramming (Fig 3D) confirming that this deletion-rescue strategy can be used to identify the domains of AF10 responsible for preserving cell identity. The AF10 mutant that cannot interact with histones phenocopied full-length AF10 (Figs 3B, D and S3B) similar to a study in human reprogramming [26]. These data imply that histone targeting may not be the primary mode of AF10 in inhibiting reprogramming.

To further investigate the crosstalk between H3K27me3 and H3K79me2, we directly compromised the enzyme complex that mediates H3K27me3, Polycomb Repressive Complex 2 (PRC2). PRC2 targets chromatin for H3K27 methylation by the enzymatic complex component EZH1/2, and promotes its spread through the EED subunit [27]. Using A395, an EED small molecule inhibitor [28], we reasoned that depletion of H3K27me3 will create more potential binding sites for AF10 which could further decrease pluripotency acquisition. A395 decreased H3K27me3 and did not affect global H3K79me2 levels (Fig 3E). Nonetheless, the deletion of AF10 increased reprogramming efficiency to the same extent irrespective of A395 addition (Fig 3F). Therefore, the histone modification specific DOT1L targeting function of AF10 is not the barrier to somatic cell reprogramming.

We next turned our attention to the two AF10 mutants that cannot interact with DOT1L (Fig 3A). There was an increase in the number of iPSCs similar to AF10 deletion when cells were rescued with AF10 mutants lacking the OMLZ domain (Fig 3D). The OMLZ deficient mutants failed to maintain H3K79me2, unlike full-length AF10 that restored almost wild-type levels (Fig 3G). Therefore, taken together with the dose-dependent increase in iPSCs upon AF10 deletion in DOT1L inhibited cells (Fig 1G), even removing only higher order H3K79 methylation can promote cell fate conversion during reprogramming.

### Higher order methylation of H3K79 preserves cell identity

To determine the impact of AF10 deletion on the degrees of H3K79 methylation, we assessed global levels of H3K79me1/2/3 during a reprogramming timecourse (Fig 4A). As compared to control cells, AF10 deletion resulted in a global reduction of both H3K79me2 and H3K79me3 to almost the same levels observed upon addition of the DOT1Li (Fig 4A-B). By contrast, AF10 deletion reduced H3K79me1 to about half of unperturbed levels (Fig 4A), with almost 5.4-fold more H3K79me1 retained as compared to DOT1Li treatment on day 4 of reprogramming (Fig 4B). Thus, the contribution of AF10 deletion to increased reprogramming efficiency is likely to occur through effects on higher order H3K79 methylations.

**Figure 4.**
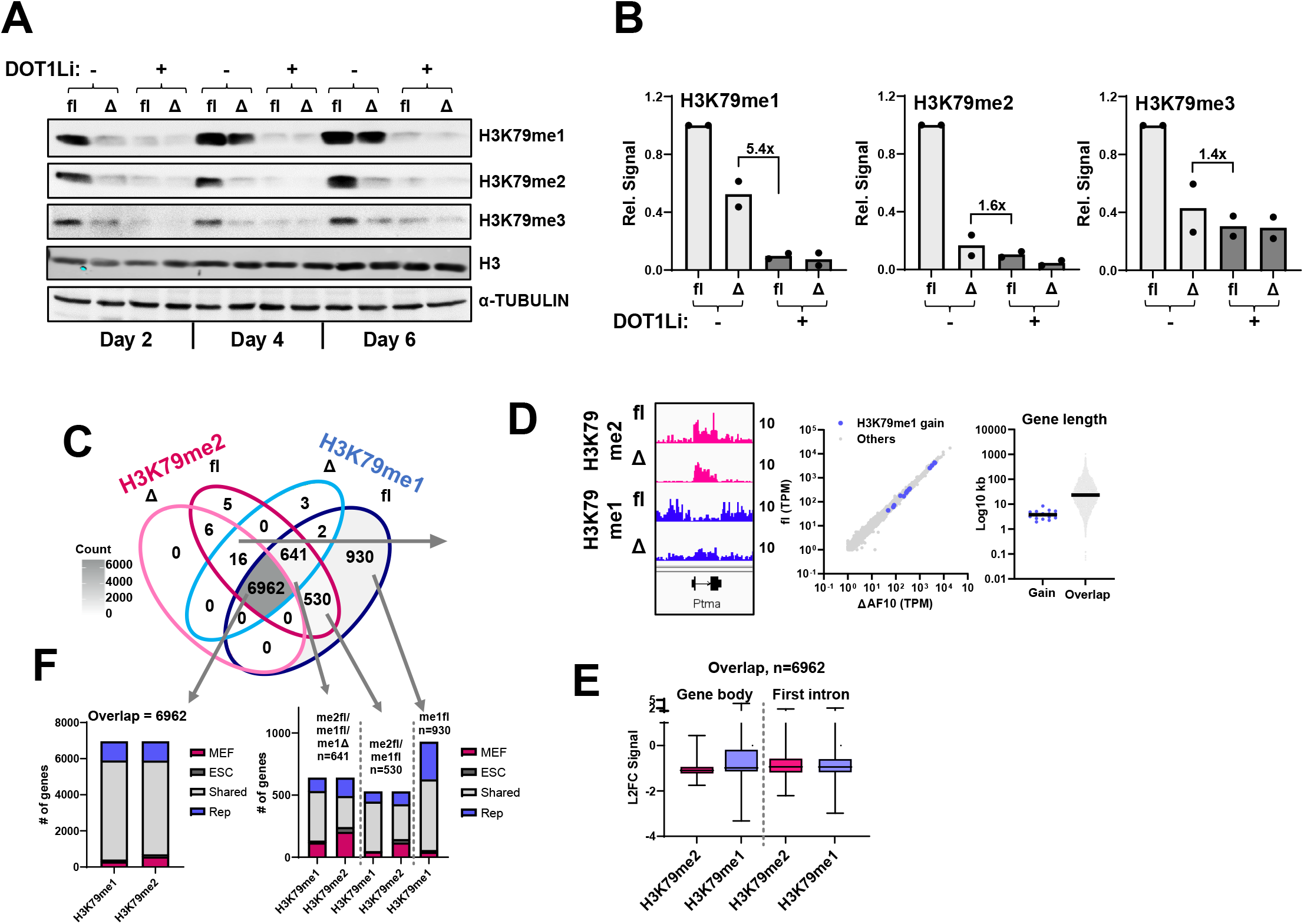
Higher order methylation of H3K79 preserves cell identity. A. Immunoblot of H3K79me1, me2, and me3 in AF10 fl or Cre deleted (Δ) conditions on days 2, 4, and 6 of reprogramming. B. H3K79me1/2/3 immunoblot signal on day 4 of reprogramming (n = 2) relative to total H3. Control treated fl condition set to 1. C. Overlap of genes with a significant H3K79me2 and/or H3K79me1 peak in AF10 wt (fl) or Cre deleted (Δ) on day 4 of reprogramming. D. Analysis of H3K79me2 unique genes that gain H3K79me1 in ΔAF10 (n=16). Left: Example IGV track. Middle: Expression (TPM) in ΔAF10 versus fl on day 4 of reprogramming. Right: Gene length compared to overlap genes with a (n=6962) H3K79me1/2 peak. E. Log2 fold change (L2FC) of normalized reads per gene body (Left) or first intron (Right) of H3K79me2 (pink) or H3K79me1 (blue) in deleted AF10 relative to fl control (Δ/f). Normalized reads are the mean (n = 2). F. MEF unique (pink), ESC unique (dark gray), MEF/ESC Shared (light gray), or Reprogramming (Rep) unique (blue) H3K79me1/2 peak status of shared (n=6962), H3K79me1 fl/K79me1 ΔAF10 /H3K79me2 fl (n=641), H3K79me1 fl/K79me2 fl (n=530), and H3K79me1 fl (n=930) gene sets.

To determine where the changes in H3K79 me1 and H3K79me2 occurred on the genome we performed quantitative ChIP-Seq with a spike in control on day 4 of reprogramming in the fl and AF10 deleted (Δ) conditions (Materials and Methods and Fig S3C). This timepoint captures the difference in H3K79me1/2 (Fig 4A-B) but precedes the formation of fully reprogrammed colonies (data not shown). Almost all the H3K79me1 and H3K79me2 peaks were found at genic locations (Fig S3D). Hence we focused on the specific genes with altered H3K79me1/me2 enrichment in an AF10 dependent manner during reprogramming. In agreement with the immunoblot, we found locations that lost both H3K79me1 and H3K79me2 (530 genes), or only H3K79me1 (930 genes), or where H3K79me2 was reduces to H3K79me1 (641 genes) in the AF10 deleted reprogramming cells (Fig 4C). Surprisingly, there were also 16 genes where H3K79me1 was gained upon AF10 deletion as H3K79me2 is reduced (Fig 4C). These genes had high levels of expression, short gene length, and efficient DOT1L activity such that nearly all H3K79me2 was converted to H3K79me1 (Fig 4D). Reduction of DOT1L catalysis via ΔAF10 enabled retention of H3K79me1 at these locations. However, these changes represented only a minority of genes, since almost 89% of H3K79me1 and H3K79me2 enriched genes were coincident in both wild-type (fl) and AF10 deleted reprogramming populations (Fig 4C). At these shared ∼7000 locations, both the H3K79me2 and H3K79me1 enrichment were reduced ∼2 fold over the entire genes body and the first intron.

Since reprogramming is the conversion from the somatic to the pluripotent state, we assessed how many of the H3K79me1/me2 genes in the fl and ΔAF10 were modified in MEFs and ESCs. We have previously shown that ∼80% of the H3K79me1/me2 locations are shared between MEFs and ESCs [29]. The vast majority of H3K79me1/2 containing genes in reprogramming populations are also modified in both ESCs and MEFs (Fig 4F). The small fraction of MEF lineage specific genes were not altered in expression levels (Fig S3E). Taken together, these results indicate that ΔAF10 in reprogramming affects H3K79 methylation levels at genes that are not ESC or MEF specific but shared in both cell types.

### Deletion of AF10 evicts higher order H3K79me at housekeeping genes

To elucidate the H3K79 methylation dynamics at these non-lineage genes, we performed k-means clustering of the ratio of H3K79me1/me2 in AF10 deleted or DOT1L inhibited samples during reprogramming compared to control (Fig 5A and S4A). We then queried how these clusters behaved in MEFs and ESCs, the start and end points of reprogramming. While the ratio of H3K79me2 remained invariant, H3K79me1 was more enriched in ESCs compared MEFs, in Cluster 1 (C1) versus C5 (Fig 5A). Interestingly, C1 genes also had high absolute levels of gene expression in both MEFs and ESCs (Fig 5A). Genes in C1-C3 tended to be smaller with a shorter first intron than those in C4 and C5 (Fig S4B). While all the clusters were comprised of genes with housekeeping functions in metabolic processes, C1 contained genes with peptide metabolism and RNA binding gene ontology categories including transcription factors, over 30 ribosomal proteins and 6 eukaryotic translation factors (Fig 5B).

**Figure 5.**
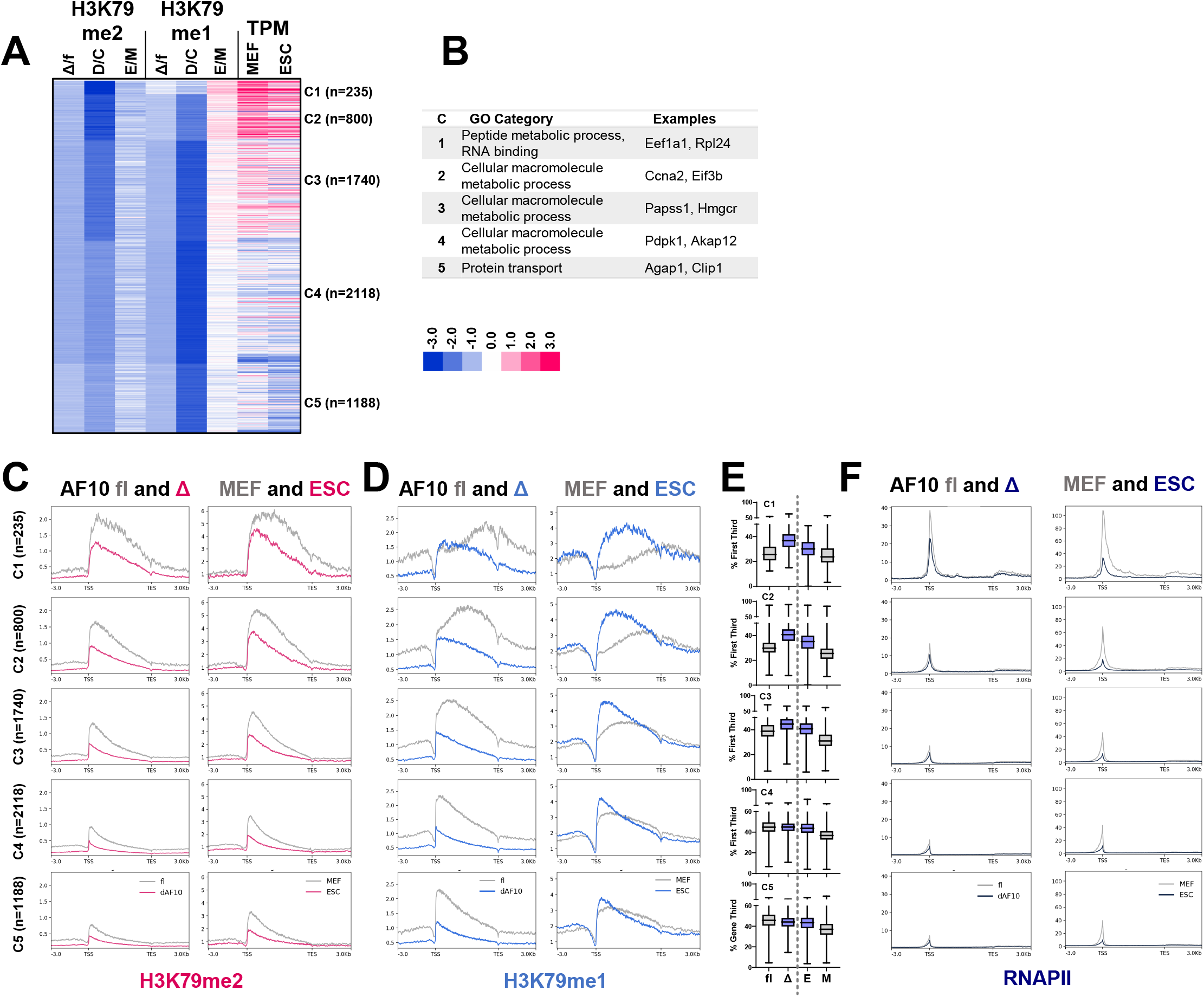
Deletion of AF10 evicts higher order H3K79me at housekeeping genes. A. Heatmap of genes with shared H3K79me1 and me2 peak on day 4 of reprogramming (Fig 4C, n=6962). Genes were clustered using the Log2 fold change (L2FC) H3K79me1 and me2 gene body enrichment in AF10-Cre relative to fl (Δ/f) and DOT1Li/Control (D/C) by k-means. L2FC H3K79me1 and me2 in ESCs relative to MEFs (E/M) and expression (TPM) was appended to the clusters. Only genes with expression information included for additional analysis (n=6081). Clusters were analyzed for (B-F): B. Gene ontology. C. H3K79me2 metaplots in AF10 wt (fl)/Cre deleted (Δ) and MEFs/ESCs. D. H3K79me1 metaplots in AF10 wt (fl)/Cre deleted (Δ) and MEFs/ESCs. E. Percent of H3K79me1 enrichment per the first third of the gene body in day 4 reprogramming control (fl) or AF10 deleted (Δ), ESCs (E), and MEFs (M). F. RNAPII metaplots in AF10 wt (fl)/Cre deleted (Δ) and MEFs/ESCs.

We next compared the pattern of enrichment of H3K79me1 and H3K79me2 upon AF10 deletion to that in MEFs and ESCs. C1 had the greatest total abundance of H3K79me2 in ESCs, MEFs, and on day 4 of reprogramming (Fig 5C) while Clusters 2 through 4 had decreasing amounts of total H3K79me2/me1 (Fig 5C-D). The deletion of AF10 broadly caused a ∼50% reduction in the H3K79me2 levels in all clusters mimicking the decrease observed between ESCs and MEFs (Fig 5C), whereas DOT1Li reduced H3K79me2 beyond what is observed in ESCs (Fig S4C). A more interesting pattern emerged in the distribution of H3K79me1 across the gene body. Similar to MEFs, in the fl control reprogramming condition, H3K79me1 is depleted at the transcriptional start site (TSS) compared to the mid-gene body, most prominently in C1 and C2 (Fig 5D). This pattern suggests that when AF10 is intact, there is almost complete conversion of H3K79me1 to H3K79me2 at the TSS. By contrast, upon ΔAF10, H3K79me1 has an apex of enrichment at the TSS (Fig 5D). This pattern resembles the H3K79me1 found in ESCs which are much more TSS-enriched for this modification compared to MEFs (Fig 5D) [29]. The distinction of TSS enrichment becomes even more obvious when reads are quantified in the proximal one third of the genes, especially in C1 and C2 (Fig 5E). Residual H3K79me1 at the TSS did not alter steady-state expression in the AF10 deleted cells (Fig S4D). Thus, deletion of AF10 during reprogramming reduces higher order H3K79 methylation and results in retention of H3K79me1 in an ESC-like pattern predominantly at highly expressed housekeeping genes.

DOT1L interacts with the phosphorylated C-terminal tail of RNA polymerase II (RNAPII) [30] and methylates genes concurrent with active transcription [3]. Therefore, we analyzed GRO-seq data, which measures rate of RNAPII transcription, from MEFs and ESCs and found that C1 genes were rapidly transcribed in both cell types (Fig S4E). C2 genes have a higher RNAPII elongation rate than the C3-C5 clusters in ESCs (C1 genes were too small for this analysis) (Fig S4F). Thus in ESCs in C1 and C2, the rapid RNAPII elongation likely does not allow DOT1L enough residence for the conversion of H3K79me1 to H3K79me2. In the AF10 deleted reprogramming conditions, the pattern of H3K79me1 in C1 and C2 resembles that in ESCs which could be due to: 1) lessened DOT1L targeting by RNAPII and/or 2) reduced DOT1L catalytic activity. We performed quantitative ChIP-seq for RNAPII on day 4 of reprogramming in fl and AF10 deleted conditions and compared to our previously generated data in MEFs and ESCs. RNAPII localization levels mirrored gene expression by cluster (Fig 5A, F) and was much higher at the TSS in MEFs as compared to ESCs. RNAPII was most highly enriched at the TSS of C1 (Fig 5F), a region that overlaps with the complete elevation of H3K79me1 to H3K79me2 in MEFs and fl reprogramming cells (Fig 5C-D). Surprisingly, deletion of AF10 reduced RNAPII at the pause site (Fig 5F), enabling the formation of the RNAPII TSS-localization pattern found in ESCs (Figs 5F and 6A). Therefore, AF10 affects H3K79me1/2 enrichment by both boosting DOT1L catalytic activity as well as regulating RNAPII engagement at the TSS. Collectively, AF10 opposes the unique H3K79me1/2 signature and RNAPII localization singular to the pluripotent fate.

**Figure 6.**
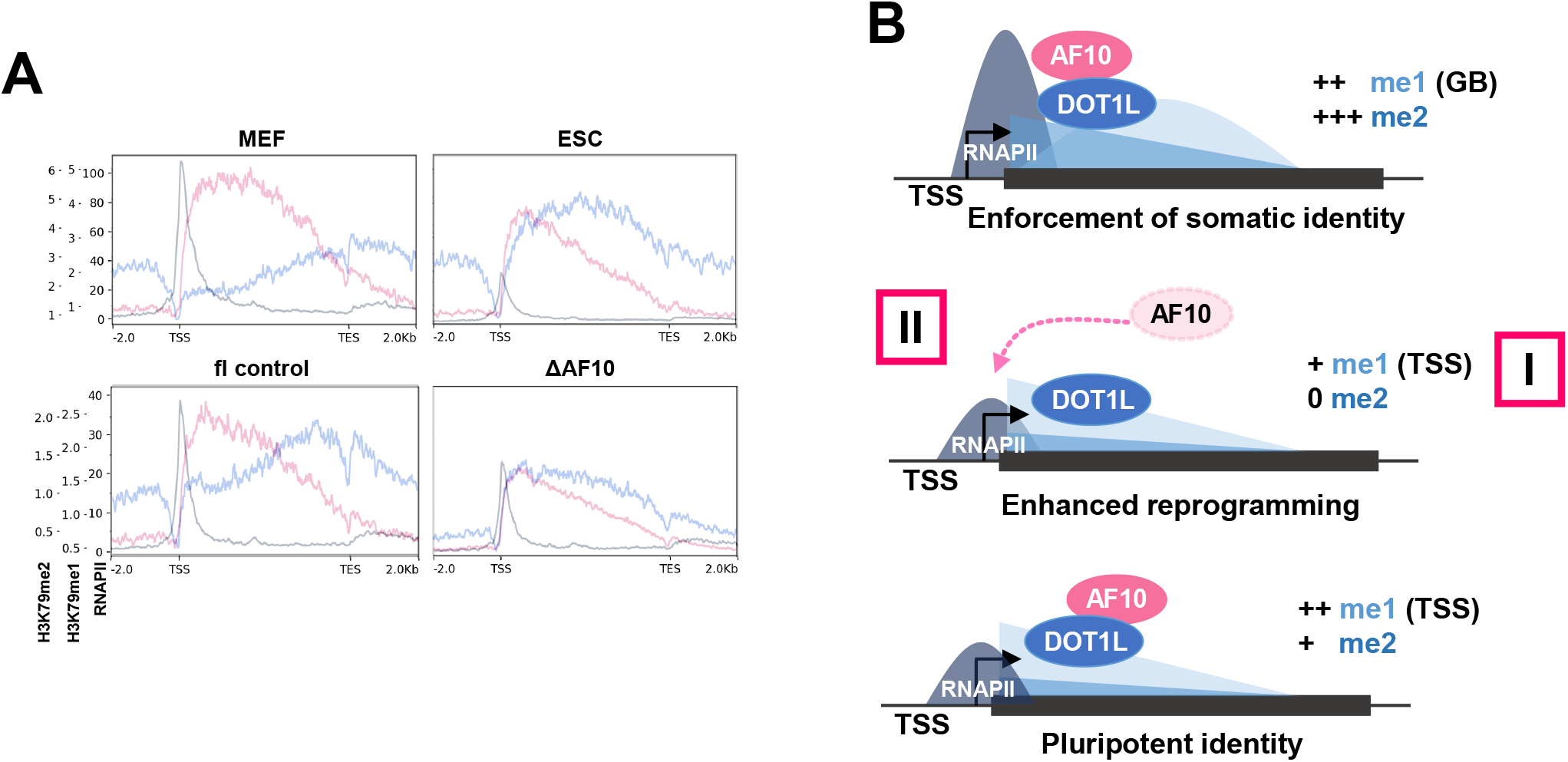
AF10 maintains cell identity by enforcing a somatic H3K79-epigenome. A. Overlapped metaplots of H3K79me1, H3K79me2, and RNAPII of genes in C1 (Fig. 5A). B. In somatic cells, AF10 and DOT1L are co-recruited to rapidly and highly expressed housekeeping genes by RNAPII yielding the maximum amount of H3K79me2. H3K79me1 is fully converted to higher order (H3K79me2/3) leading to depletion of H3K79me1 at the TSS. In ESCs, the relatively reduced TSS-associated RNAPII leads to lower H3K79me2 and a TSS apex of H3K79me1. ESCs have increased transcription of housekeeping genes suggesting H3K79me2 functions in a negative feedback loop. Deletion of AF10 during reprogramming reduces DOT1L catalytic function resulting in retained H3K79me1 where H3K79me2 is normally highly enriched. AF10 deletion also lessens RNAPII at the TSS. Thus, acquisition of the pluripotent fate is increased by: I. Lowering the total levels of H3K79me and/or II. Establishing an ESC-like TSS pattern of H3K79me1 deposition and RNAPII engagement.

Given that AF10 seems to have a negative effect on RNAPII enrichment at the TSS during reprogramming we wondered why there was an increase in expression of AF10 in pluripotent cells (Figs 1A and S1A). There is an alternatively spliced isoform of AF10 which uses an internal ATG that skips the first ∼80 amino acids that encode zinc fingers that help mediate nucleosomal interaction [9,31,32]. Interestingly, there is an increase in expression of this shorter isoform during reprogramming (Fig S4G) which may act as a dominant negative partner to sequester DOT1L in ESCs.

## DISCUSSION

Taken together our results favor a model in which DOT1L may initially be targeted to loci by RNAPII (Fig 6B), the AF10 interaction is essential for maintaining or stabilizing RNAPII at the promoter. The presence of AF10 increases higher order H3K79 methylation which is unfavorable for pluripotency. Even without the complete loss of H3K79 methylation (Fig 4A), merely reducing H3K79me2 and/or reorganizing the enrichment pattern of H3K79me1 with AF10 deletion (Fig 6, I vs. II) can cause a remarkable 4-fold gain in reprogrammed colonies (Fig 1E). These changes occur especially at highly expressed housekeeping genes with a rapid rate of transcription (Figs 5 and S4). Compared to somatic cells, ESCs have a lower TSS-enrichment of RNAPII (Fig 5F) leading to reduced global levels of H3K79me2 and preservation of H3K79me1 at the TSS (Figs 5C-D and 6A). This is mirrored in AF10 deleted reprogramming cells, even though they are a heterogenous population, where lowered RNAPII at the TSS generates a high H3K79me1-low H3K79me2 pattern at highly transcribed genes.

The pluripotent-specific enrichment and pattern of H3K79me1/2 is not only a consequence of reduced RNAPII localization at the TSS in ESCs. Pluripotent stem cells express an alternative compromised isoform of AF10 (Fig S4G), and exclude DOT1L from the nucleus [15] to actively maintain low levels of H3K79me2. Combined, the unique epigenetic pattern and RNAPII profile may mitigate negative transcriptional feedback (Fig 6B) [29]. ESCs have a much greater need for biosynthetic output than somatic cells [33,34] and are likely to regulate transcription of housekeeping gene in a distinct manner [29]. AF10 mediated enhancement of DOT1L catalytic activity during reprogramming detains RNAPII enrichment at the TSS, similar to the pattern found in MEFs (Fig 6). Such a function for AF10 may to be evolutionarily conserved since a greater enrichment and degree of H3K79me has been shown to negatively impact transcriptional rate in *C. elegans* [35]. Specifically, the presence of the AF10 homolog ZFP-1 inhibits the transition of RNAPII pausing to productive elongation [35].

Although AF10 greatly enhances pluripotency acquisition (Fig 1), its deletion results in few transcriptional changes (Fig 2). This result mirrors our recent report that showed DOT1Li yields few significant transcriptional alterations despite global eviction of H3K79me, and furthermore, none of differentially expressed genes are causal for enhanced reprogramming efficiency [18]. AF10 is in the same pathway as DOT1L in human fibroblast reprograming indicating it is a broad regulator of cell fate. However, its deletion enhanced reprogramming by downregulating fibroblast gene expression and upregulating the pluripotency network in the human system [26]. Previous studies have shown that abrogation of AF9 [11] and AF10 [10] dramatically decrease H3K79me at target genes, yet the transcriptional effect is modest. It is interesting to note that AF10 deletion tempers the already modest DOT1Li effect on steady state mRNA (Fig 2), indicating that they are in the same pathway. Therefore, the majority of steady state transcriptional alterations do not contribute to pluripotency acquisition, and instead AF10 functions as an epigenome modifier to mediate higher order H3K79 methylation to maintain cell identity.

We do not find any evidence that H3K79me2 globally opposes H3K27me3 in reprogramming to iPSCs (Fig 3E), although direct local effects cannot be excluded. Inhibition of EED and loss of H3K27me3 did not increase H3K79me2 or inhibit reprogramming (Fig 3E-F). A recent study in leukemia found reduction of H3K27me3 at genes where H3K79me2 is removed with lower, but not higher, concentrations of EPZ-5676 [36]. Perhaps this reduced concentration of EPZ-5676 mimics DOT1L attenuation via ΔAF10 in our study. DOT1L knockout in lymphocytes downregulated EZH2 and de-repressed genes normally shut off by H3K27me3 [37,38]. Inhibition of DOT1L and EZH2 have both shown promise in pre-clinical MLL-R studies suggesting that H3K79me and H3K27me3 are functionally connected in leukemia [39].

We find that AF10 maintains cell identity by boosting DOT1L catalytic activity, but not through the H3K27 interaction domain (Fig 3). In contrast, DOT1L targeting is important in leukemias where AF10, AF9 and ENL are some of the most frequent fusion partners in rearrangements of the mixed-lineage leukemia (MLL) gene. In these MLL-rearranged (MLL-R) leukemias, DOT1L is recruited to MLL target loci with consequent aberrant H3K79me deposition [40–45]. Depletion of AF9 and ENL indicates that they do not function in the H3K79me2-mediated maintenance of cell identity (Fig 1B). We did not observe an increase in cell death upon AF9 or ENL depletion (data not shown) but further studies can determine whether the DOT1L- or transcriptional complex-interaction functions of AF9 and ENL contribute to their phenotype in reprogramming.

Taken together our studies have uncovered a mechanism by which changes in the epigenetic distribution of a gene body associated histone modification can cause a profound change in cell identity.

## MATERIALS AND METHODS

### Mice and breeding

Mice were maintained in agreement with our UW-Madison IACUC approved protocol. Reprogramming mice are homozygous for the Oct4-2A-Klf4-2A-IRES-Sox2-2A-c-Myc (OKSM) transgene at the *Col1a1* locus and either heterozygous or homozygous for the reverse tetracycline transactivator (rtTA) allele at the *Rosa26* locus [14]. Conditional AF10 knockout mice with LoxP sites before exon 17 and after exon 18 (OM-LZ domain) were a kind gift from Dr. Aniruddha Deshpande (Sanford Burnham Prebys Medical Discovery Institute) and Cre-mediated deletion results in loss of AF10 protein [10]. These mice were bred to the reprogrammable mouse strain to generate strains homozygous for flox (fl)-AF10, homozygous for OKSM, and either heterozygous or homozygous for rtTA. DR4 mice are genetically resistant to geneticin (G418), puromycin, hygromycin, and 6-thioguanine and acquired from Jackson labs.

### Cells and culture

MEFs were isolated from embryos at day E13.5 of time mated mice as previously described [46]. MEFs were maintained in MEF media (DMEM, 10% FBS, 1x non-essential amino acids, 1x glutamax, 1x penicillin/streptomycin, and 2-Mercaptoethanol - 4 μl/500ml). DR4 feeder MEFs were expanded for 3 passages and irradiated with 9000 rad. iPSCs were picked from reprogrammed wells, genotyped, and passaged at least 5 times without doxycycline. V6.5 ESCs and iPSCs were grown in ESC media (knock-out DMEM, 15% FBS, 1x non-essential amino acids, 1x glutamax, 1x penicillin/streptomycin, 2-Mercaptoethanol - 4 μl/525ml, and leukemia inhibitory factor) on plates coated with 0.1% porcine gelatin and DR4 feeder MEFs. Platinum-E (PLAT-E) are 293T cells engineered to express retroviral packaging components Gag, Pol, and Env, and are maintained in DMEM, 10% FBS, 1x penicillin/streptomycin, 10 μg/mL puromycin, and 1 μg/mL blasticidin.

### Genotyping

Genotyping of mouse litters was performed on ear clippings at 21 days. Genotyping of reprogramming experiments was performed 2 days post Cre recombinase treatment. Tissue or cells were digested in 100 mM Tris-HCl pH 8.5, 5 mM EDTA, 0.2% SDS, and 200 mM NaCl with 20 μg Proteinase K for 2 hours at 65°C, followed by DNA precipitation with an equal volume of isopropanol and 20 μg glycogen. AF10 genotyping was performed with: 5’-CACAGCCTACTTCAAAGAAC-3’, 5’-TAGTCATGGGAATGGAGATG-3’, and 5’-ATTAGAGTCCATCCCACTTC-3’ primers..

### Reprogramming

Cells were plated in MEF media on glass coverslips (Warner Instruments, 64-0714) that had been coated with 0.1% porcine gelatin, at a density of 30,000-50,000 cells per 12-well or 10,000-20,000 per 24 well. Reprogramming (OSKM expression) was initiated with 2 μg/mL doxycycline hyclate (Sigma-Aldrich, D9891). Reprogramming cells were treated with vehicle (DMSO) control, SGC0946 (ApexBio, A4167), EPZ-5676 (MedChem Express, HY-15593), or EPZ004777 (MedChem Express, HY-15227) beginning at day 0 of reprogramming, at 5, 2.5, 1, 0.2 μM, as indicated. SGC0946 was used at 5 μM in all additional reprogramming experiments. Cells were treated with 1 uM EED inhibitor A-395 hydrochloride (Millipore Sigma, SML1923) beginning on day 1 of reprogramming. Gamma-irradiated feeder DR4 MEFs were added at 0.5x confluency by day 2 of reprogramming. Media was switched to ESC media by day 2 of reprogramming. Media with fresh doxycycline and drugs was replenished every two days of reprogramming. In the case of AF10 deletion experiments, Cre recombinase was added 1-2 days before doxycycline, or at the indicated timepoints (Methods, Viral transduction).

In the case of siRNA depletion experiments, cells were plated 1 day post doxycycline treatment and transfected with 20 nmol siRNA overnight. Media was switched to ESC media containing knock-out serum replacement (KSR) and feeder MEFs were added on day 2. Cells were then transfected with 40 nmol siRNA on days 3, 5, and 7 to account for increased cell number.

Stable colony formation was measured by removing media containing drugs and doxycycline, washing cells with ESC media, and replacing with ESC media free of doxycycline and drugs. Coverslips were fixed 2-4 days post doxycycline removal and assessed for sustained NANOG expression. Reprogramming timing was carefully adjusted for individual MEFs such that transgenes were activated long enough to produce *bona fide* colonies, but not so long that counting NANOG positive colonies in the presence of doxycycline was impossible due to overcrowding.

### Reprogramming statistical analysis

Reprogrammed colonies were quantitated using fluoroscopy and were comprised of a grouping of at least 4 NANOG+ cells. At least 3 biological replicates were performed on separate days and/or with a separate MEF isolation. Biological replicate number (n) is included in all legends. Biological replicates consisted of at least 2 technical replicate coverslips per condition. The number of colonies in the control condition (fl + vehicle control treatment) was averaged per biological replicate, and set to 1 (or as specified in legends). The fold change in reprogrammed colonies per technical replicate was calculated against the control average. Reprogramming efficiency was plotted in Graphpad Prism and statistically analyzed in R using the ggplot2, Rmisc, car, emmeans, PMCMRplus packages. The Shapiro-Wilk test was used to test normality and the Fligner-Killeen tests was used to assess homogeneity of variances. If both conditions were met, data were analyzed using one-way ANOVA with the Tukey-Kramer test post-hoc to make pairwise comparisons. If the homogeneity of variances was not met, Welch’s ANOVA with post-hoc pairwise t-tests using the Welch adjustment and the Holm method to correct multiplicity of pairs, was employed. All statistical testing and p-values explained in the legends.

### siRNA transfection

siRNA was diluted in up to 50 μL serum-free DMEM per 12 well. 1 μL of DharmaFECT was incubated with 49 μL of serum-free DMEM per 12 well, for 5 minutes. The transfection reagent was combined with the diluted siRNA and incubated at room temperature for 20 minutes. Cells were transfected dropwise. The following siRNAs were used in reprogramming experiments and purchased from Dharmacon (horizon): siDot1l (J-057964-12), siMllt10 (J-042898-09), siMllt1 (D-063284-01), siMllt3 (D-050750-03), non-targeting control (D-001810-01).

### Viral transduction

Cre recombinase (Ad5-CMV-Cre) and empty control (Ad5-CMV-Empty) purified adenoviruses were purchased from the Vector Development Lab (Baylor College of Medicine). 2.4×10^9^ pfu of virus was resuspended in 5 mL of serum-free DMEM and added to 1-2 million MEFs in a 10-cm dish. Virus containing media was removed after 1 hour and replaced with MEF media. Reprogramming was initiated 2 days post-transduction.

AF10 domain mutant retroviruses were packaged in PLAT-E cells. Cells were treated with 10 μg/mL puromycin and 1 μg/mL blasticidin for 4 days to select for cells expressing packing components. Media was changed to antibiotic free DMEM+10% FBS on 15-cm dishes at 60% confluency 2 hours before transfection. 15 μg DNA was incubated in 600 μL Opti-MEM and 60 μL of 1 mg/mL linear PEI (Polyscience, 23966-2) for 15 minutes at room temperature before adding to cells dropwise. Media was changed to MEF media plus 20 mM HEPES 4 hours post-transfection. Virus-containing media was collected 48- and 72-hours post-transfection, filtered through a 0.45 um PVDF filter, and added directly to MEFs or frozen at −80°C for long term storage. MEFs were infected three times with the following strategy: 4 hours of infection (50% viral media, 50% fresh MEF media, 10 μg/mL Hexadimethrine Bromide), 4 hours of rest (MEF media), overnight infection, 4 hours of rest, 4 hours of infection.

### Immunofluorescence

Cells were fixed for 10 minutes at room temperature in 4% paraformaldehyde/1xPBS. Cells were permeabilized in 0.5% Trition-X/1xPBS and washed in wash buffer (0.2% Tween-20/1xPBS), each for 10 minutes. Coverslips were blocked for 30 min in blocking buffer (5% normal goat serum, 0.2% Tween-20, 0.2% fish skin gelatin, 1x PBS) and probed for 1 hour in blocking buffer containing α-NANOG (1:100; Cosmo Bio USA, REC-RCAB001P or 1:1000; Cell Signaling Technology, 8822S) or α-OCT4 (1:400; Cell Signaling Technology, 2840) on parafilm. Coverslips were washed twice in wash buffer and then probed with α-Rabbit IgG-Dylight 488 (1:1000; ThermoFisher, 35552) for 1 hour. Coverslips were washed once, incubated with 4′,6-Diamidino-2-phenylindole dihydrochloride (1:10,000; Millipore Sigma, D8417) in wash buffer, and then washed a second time before mounting with Aqua-Poly/Mount (Fisher Scientific, NC9439247).

### RT-qPCR

RNA was isolated from cells with Isolate II RNA Mini Kit according to the manufacturer’s instructions (Bioline, BIO-52702). 1 μg of RNA was converted to cDNA using qScript (Quanta, 95047). Expression was measured with 20 ng of cDNA (calculated based on starting RNA quantity) in 10 μL reactions with SYBR Green (Bio-Rad, 1725124). Relative expression for knock-down was calculated using the Delta-Delta Ct method with *Gapdh* as the control reference, and non-targeted siRNA condition set to 1. Relative expression AF10 isoforms was calculated using Delta Ct method with *Gapdh* as the control reference. The following primers were used:

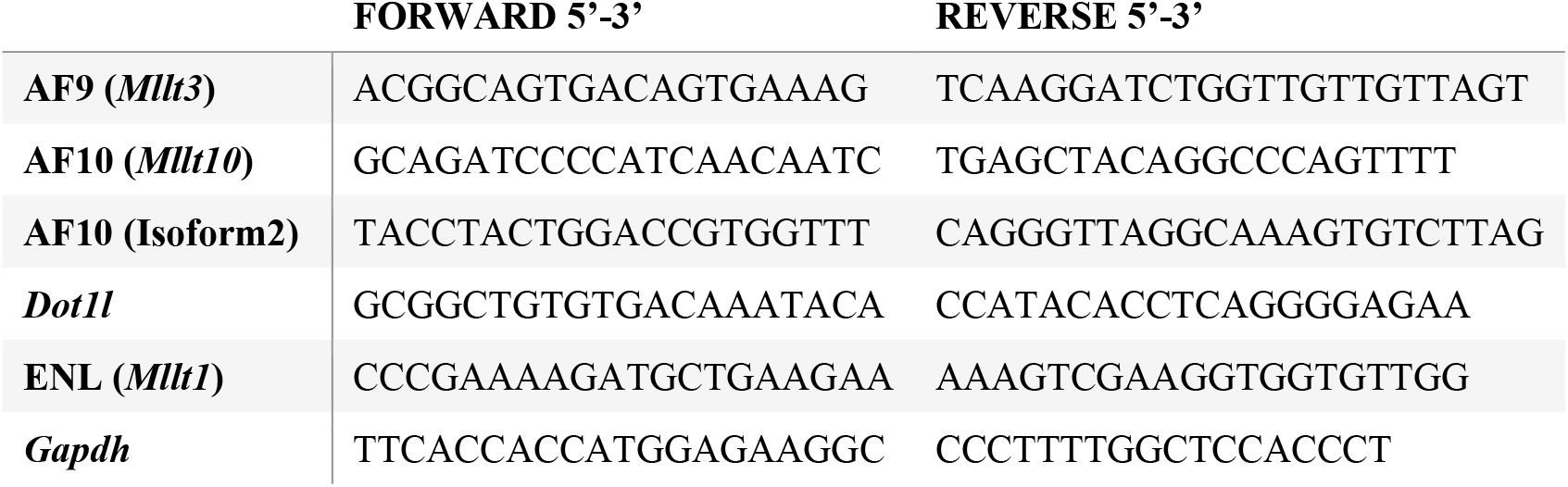

### Immunoblot

Approximately 1 million whole cells were lysed in SUMO buffer - a fresh mixture of ¼ part I (5% SDS, 0.15 M Tris-HCl pH 6.8, 30% glycerol), ¾ part II (25 mM Tris-HCl pH 8.3, 50 mM NaCl, 0.5% NP-40, 0.5% deoxycholate, 0.1% SDS), and 1x cOmplete protease inhibitors (Roche, 4693132001) - and sonicated with a microtip at 20% amplitude for 5 seconds on ice. Protein concentration was measured with DC Protein Assay kit II (BioRad, 5000112) against a BSA standard curve and measured with Synergy H4 microplate reader. 20-25 μg of protein was loaded on H3K79me1/2 gels, 10 μg on H3K27me3 and H3K79me3, and 15 μg on FLAG-AF10 gels. Blots were transferred to nitrocellulose membranes, and blocked and probed in 5% milk, PBS, 0.1% Tween-20. The following antibodies were used: FLAG-M2 (1:5000; Sigma-Aldrich, F3165), H3K79me1 (1:1000; Diagenode, C15410082), H3K79me2 (1:1000; Active Motif, 39143), H3K79me3 (1:1000; Diagenode, C15410068, or 1:1000; Abcam, ab2621), H3K27me3 (1:1000; Cell Signaling, 9733S), total Histone H3 (1:3000; Cell Signaling, 3638S) and α-TUBULIN (1:3000; Cell Signaling, 3873). Immunoblot images were captured with ECL on an ImageQuant LAS 4000 or with LI-COR Odyssey. Tiff files were quantitated with Image Studio Lite V5.2 using the Add Rectangle function to calculate signal and background intensities.

### Plasmids

AF10 domain mutants in the MSCV_IRES_YFP retrovirus plasmid were provided by Dr. Aniruddha Deshpande (Sanford Burnham Prebys Medical Discovery Institute). The full length (FL) vector contains the human AF10 cDNA isoform X6 (XM_024448184.1), NTD-OMLZ (NO) encodes aa 1-724, ΔOMLZ (ΔO) encodes aa 1-730, 798-1084 with the OMLZ domain replaced by a methionine and histidine, and NLS-CTD encodes aa 207-1084. 3xFLAG was inserted N-terminal of AF10 in all constructs between the EcoRI and BamHI sites to facilitate immunoblot detection.

### Flow cytometry sorting

AF10 domain mutant (MSCV_AF10_IRES_YFP) containing cells were FACS sorted before reprogramming. Cells were trypsinized, washed in PBS, resuspended in MEF media, and filtered through cell strainer capped tubes (Corning, 352235) before collecting the 530^+^ population on a BD FACS Aria II (UW Carbone Cancer Center, Grant #: 1S10RR025483-01). Uninfected MEFs were used to set the gates (Fig S3A). Flow cytometry images were generated in FlowJo.

### RNA-Seq

RNA was isolated from 2 independent biological replicate reprogramming experiments starting from independent MEF isolations, using 1 mL of TRIzol. After 5 minutes of incubation, 200 μL of chloroform was added and mixed thoroughly. The aqueous phase was isolated after spinning the samples at 21xg for 15 min at 4°C. 0.53 volumes of ethanol was added, mixed, and spun through Qiagen RNeasy column at 21xg for 30 seconds. DNA was digested on the column with 10 μL RNAse free DNase mixed with 70 μL RDD buffer (Qiagen, 79254) for 30 min at room temperature. The RNA was then washed with 500 μL of buffer RW1 and RPE, each. RNA was eluted with 30 μL of water. 2 μg of RNA was used as starting material for library construction with TruSeq RNA Sample Prep Kit V2 (Illumina, RS-122-2002) following the manufacturer’s instructions. Libraries were assessed with Qubit dsDNA HS assay and bioanalyzer3.0 before sequencing on HiSeq 2500 (UW-Madison Biotechnology) single end, 100 bp. Samples had 20-30 million reads.

### RNA-Seq analysis

Sequencing quality was assessed with FastQC (http://www.bioinformatics.babraham.ac.uk/projects/fastqc/). Reads were processed before alignment with the FASTX-toolkit (http://hannonlab.cshl.edu/fastx_toolkit/). First low-quality reads were removed with the following parameters: fastq_quality_trimmer -Q33 -t 28 - l 20. Adapters were removed with the following parameters: fastx_clipper -Q33 -a adapter_sequence. The 5’ end of the read (length determined by FastQC per base sequence content) was removed with the following parameters: fastx_trimmer -Q33 -f 11. Reads were then aligned to mm9 with an allowance of up to 2 mismatches using RSEM [47] with the following parameters: rsem-calculate-expression -bowtie-m 200 --bowtie-n 2 --forward-prob 0.5 --seed-length 28. DE gene analysis was initiated by generating matrices of the “expected counts” from RSEM .gene.results files of the two replicate samples for DE comparison on R. These matrices were analyzed using EBSeq [48] with the follow parameters: MedianNorm, 5 iterations, target FDR = 0.05, and posterior probability of being DE (PPDE) > 0.95 on R. Genes were considered DE if the posterior FC (PostFC) was greater than or equal to 2 (upregulated) or less than or equal to 0.5 (downregulated).

Expression changes were assessed with TPM values collected in a matrix across all samples from RSEM .gene.results files using R. TPM values of DE genes from all sample comparisons were assessed with a Pearson correlation and displayed using the R package “pheatmap”. To cluster DE genes, TPM values from replicates were averaged, 1 was added, and then they were calculated as Log2 fold change relative to the MEF control sample (fl). The Log2 FC values were clustered using Gene Cluster 3.0 [49] with the following parameters: Organize genes, 100 runs, k-means, Euclidean distance. Various numbers of clusters were tested, of which 7 were chosen based on downstream analysis. Clustered gene Log2FC values were displayed using Java TreeView [50]. Clusters were analyzed using DAVID gene ontology [51] with the following parameters: Gene Ontology (GOTERM_BP_4, GOTERM_MF_4), Functional annotation clustering. Categories with a p-value less than 5.0E-3, in cluster(s) with the highest enrichment score were displayed. Motif analysis was performed with HOMER [52] with the following parameters: findMotifs.pl -start −1000 -end 100 -len 6,10.

### ChIP

Reprogrammable AF10fl MEFs that had been treated with CRE-recombinase or control virus (day - 2) were plated on day 0 of reprogramming at 2 million cells per 15 cm dish in ESC media with feeder MEFs (Methods, Reprogramming). Media was refreshed on day 2, and cells were fixed on day 4 of reprogramming. Cells were fixed for 10 minutes with 1% formaldehyde, in solution, rotating at room temperature. Fixing was quenched with 0.14 M glycine for 5 minutes, and the cells were spun down at 300xg for 3 minutes. Cells were washed 3 times with PBS and flash frozen. 25-30 million cells were resuspended in 1 mL lysis buffer (1% SDS, 50 mM Tris-HCl pH 8, 20 mM EDTA, 1x cOmplete (Roche, 4693132001) protease inhibitor), and sonicated on a Covaris S220 Focused-ultrasonicator with the following parameters: 21 cycles of 45 seconds ON (peak 170, duty factor 5, cycles/burst 200), 45 seconds OFF (rest) in 6-8°C degassed water. 25 μL samples were taken before (0 cycles), at the mid-point, and at the end to test sonication fragmentation. Sonication test samples were resuspended in 3 volumes of water with 10 μg of RNaseA and incubated at 37°C for 30 minutes. 20 μg of Proteinase K was added, and crosslinks were reversed overnight at 65°C. DNA was cleaned and isolated with phenol-chloroform extraction using phase-lock tubes (VWR, 10847-802). 5 μg was run on a 1.5% agarose gel to ensure fragments of 200-400 bp were generated. Sonicated DNA was quantified with Qubit dsDNA HS (ThermoFisher Scientific, Q32854), aliquoted, and stored at −80°C.

The H3K79me1/2 immunoprecipitation was set up with 7.1 ug of reprogramming day 4 chromatin plus 1:53 (133 ng) of human 293T spike in chromatin for normalization [53], generated as above (Fig S3C). The RNAPII immunoprecipitation was set up with 13 ug of reprogramming day 4 chromatin plus 1:53 (247 ng) of human 293T spike-in chromatin. Chromatin was diluted 1:10 with dilution buffer (16.7 mM Tris-HCl pH 8, 0.01% SDS, 1.1% Trition-X, 1.2 mM EDTA, and 167 mM NaCl), and immunoprecipitated with 5 ug H3K79me1 (Abcam, ab2886, Lot GR274715-4), 5 ug H3K79me2 (Active Motif, 39143, Lot 16918002), or 10 ul of RPB1 NTD D8L4Y (Cell Signaling Technology, 14958S) antibody for 16 hours, rotating at 4°C. 25 μL of Protein A (ThermoFisher Scientific, 10002D) and 25 μL of Protein G (ThermoFisher Scientific, 10004D) Dynabeads that were washed once in PBS, 0.02% Tween-20 and once in dilution buffer, were added. Chromatin was incubated, rotating, with beads for 2 hours at 4°C. Beads were washed twice, rotating at 4°C for 5 minutes, with 1 mL of each of the following buffers: Low salt (50 mM HEPES pH 7.9, 0.1% SDS, 1% Triton X-100, 0.1% Deoxycholate, 1 mM EDTA pH 8.0, 140 mM NaCl), High salt (50 mM HEPES pH 7.9, 0.1% SDS, 1% Triton X-100, 0.1% Deoxycholate, 1 mM EDTA pH 8.0, 500 mM NaCl), LiCl (20 mM Tris-HCl pH 8, 0.5% NP-40, 0.5% Deoxycholate, 1 mM EDTA pH 8.0, 250 mM LiCl), and TE (10 mM Tris-HCl pH 8, 1 mM EDTA pH 8) using a magnetic rack. Beads were incubated with 250 μL elution buffer (50 mM Tris-HCl pH 8, 1 mM EDTA pH 8, 1% SDS) for 10 minutes at 65°C, shaking. 200 μL of TE with 0.67% SDS was added, and incubated shaking at 65°C for 10 minutes. 10 μg RNase A was added and incubated at 37°C for 30 minutes. 40 μg of Proteinase K was added, and incubated for 12-16 hours at 65°C to reverse crosslinks. DNA was cleaned with phenol-chloroform phase separation with phase lock tubes (VWR, 10847-802). DNA was precipitated with 1 volume isopropanol, 1/10^th^ volume of 3M NaActate, and 20 μg glycogen, and washed with 70% ethanol. DNA was dissolved in 30 μL ultrapure water and concentrated to 5 μL using a SpeedVac for library preparation.

The entire ChIP sample was used as starting material for library construction with Ovation Ultralow System V2 using reagents at 0.5X and adapters diluted 1/5 in water, following the manufacturer’s instructions (NuGEN, 0344). The libraries were separated on an 1.5% agarose gel, cut from 200-400 bp for an additional size selection step. The gel fragment was purified using the MinElute Gel Extraction Kit (Qiagen, 28604). Libraries were assessed with Qubit quantitation and Bioanalyzer3.0. Single-end 50 bp sequencing was performed on a HiSeq 4000 for a minimum of 20 million reads (NUSeq Core Northerwestern Medicine, Center for Genomic Medicine). See Table 1 for sequencing depth, antibody, and normalization details.

**Table 1.**
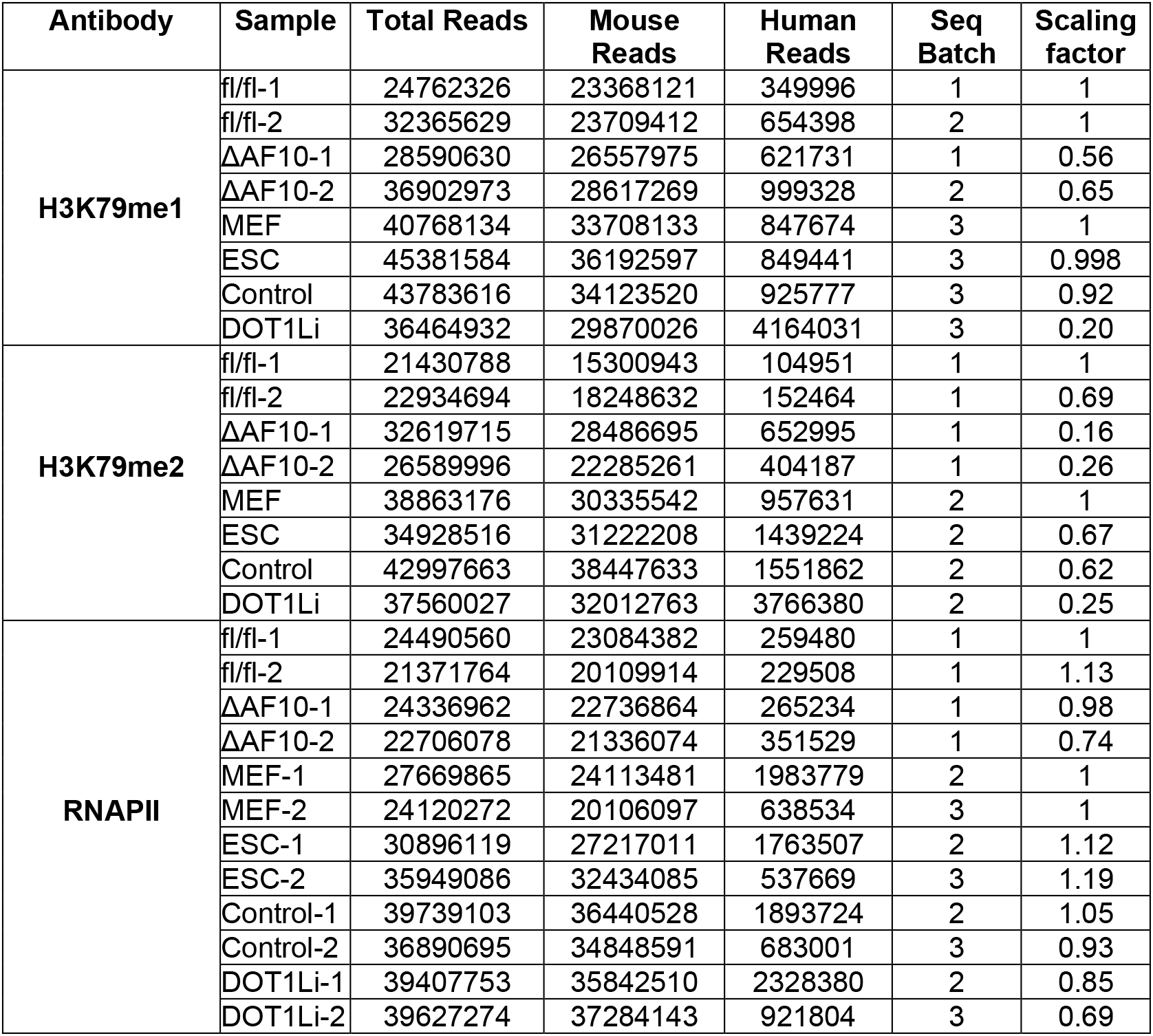
ChIP-Seq sample information.

### ChIP analysis

Ambiguous reads (that could align to both the mouse and human genomes) were “pre-cleared” by aligning first to the human hg19 genome with Bowtie2 [54]. Samtools-1.2 [55] was used to convert .sam files into .bam files using “view” and, and then sort .bam files with the “sort” command. Sample reads that did not align to the human genome were isolated from sorted bam files with Sam-1.2 “view -f4”, converted to fastq files with “bam2fq”, and then aligned to mm9 genome as above (Fig S5A). The opposite pipeline was used to calculate the number of reads that uniquely aligned to the human genome (Fig S5A), which were used for spike in normalization (see Table 1). Peaks were called with MACS2 [56,57] with the following parameters: H3K29me2 --broad -p 0.0001 and H3K79me1 –broad -p 0.001. Peaks were annotated with HOMER [52] to the mm9 genome using annotatePeaks.pl. High confidence genes were identified by isolating genes with H3K79me peaks within gene bodies (5’UTR, exon, intron, 3’UTR, and/or TTS) for each ChIP-Seq replicate, and then taking the consensus using Venny2.0 (https://bioinfogp.cnb.csic.es/tools/venny/).

The clustered heatmap (Fig 5D) was created by calculating H3K79me enrichment per gene body with BEDTools2.0 [58] multibamCov. Reads were normalized per scaling factor (see Table 1), and then per kilobase of gene length. The two ChIP-seq replicate normalized quantitations were averaged per gene and the Log2 fold change (L2FC) was calculated per indicated conditions. All box and whiskers plots were generated in Graphpad Prism as: the box limits - 25th to 75th percentiles, the center line - median, and the whiskers - minimum and maximum values. Clustering was performed with H3K79me1 and me2 L2FC in ΔAF10/fl and DOT1Li/Control using Gene Cluster 3.0 [49] with the following parameters: Organize genes, 100 runs, k-means, Euclidean distance. Various numbers of clusters were tested, of which 4 were chosen based on downstream analysis. The expression measured as transcripts per million (displayed on the heatmap as Log2 (TPM divided by 25)) and L2FC H3K79me1 and me2 in ESCs relative to MEFs were matched back to the clustered list. Clusters were displayed with Java TreeView [50]. Clusters were analyzed using DAVID gene ontology [51] with the following parameters: Gene Ontology (GOTERM_BP_4, GOTERM_MF_4), Functional annotation clustering. Categories with a p-value less than 5.0E-3, in cluster(s) with the highest enrichment score were considered.

Deeptools [59] was used to generate metaplots. First, a bigwig file normalized per human spike-in, that was the average of the two sorted .bam ChIP-Seq replicates was created with the following parameters: bamCompare --scaleFactors (see Table 1) --operation mean --binSize 10 - -ignoreForNormalization chrX. A matrix of reads per cluster was generated with the Deeptools computeMatrix command. Profiles were created with: scale-regions -b 3000 -a 3000 -- regionBodyLength 5000, or as listed. Deeptools was used to compare ChIP-Seq replicates using a Pearson correlation of normalized bigwig files. A matrix of enrichment per 1 kb bins over the genome was made with: multiBigwigSummary -bs 1000, which was displayed with: plotCorrelation --corMethod pearson --skipZeros --removeOutliers. IGV was used to visualize ChIP-Seq tracks of normalized bigwig files.

### Published datasets

ESC RNA-Seq data and bulk expression of DOT1L and associated factors was assessed from data previously generated [18] and can be obtained at NCBI GEO GSE160580. Single cell expression and co-expression of DOT1L and associated factors were analyzed as described [20] and can be obtained at NCBI GEO GSE108222. MEF, ESC, and day 4 of reprogramming in Control or DOT1Li H3K79me1/2 and RNAPII ChIP-seq datasets were processed as described in Methods, ChIP-seq analysis, and can be obtained at NCBI GEO GSE190391. GRO-Seq data was analyzed from GSE27037 [60]. Transcription elongation rates were derived from Figure 3, Source Data 1 [5].

## DATA AVAILABILITY

The datasets produced in this study are available in the following databases:

- RNA-Seq data: Gene Expression Omnibus GSE214139 (https://www.ncbi.nlm.nih.gov/geo/query/acc.cgi?acc=GSE214139).
- ChIP-Seq data: Gene Expression Omnibus GSE214141 (https://www.ncbi.nlm.nih.gov/geo/query/acc.cgi?acc=GSE214141).

## ACKNOWLEDGEMENTS

This work was supported by a UW-Madison Stem Cell and Regenerative Medicine Center postdoctoral award to CKW, and NIH-NIGMS R01GM113033 to the RS lab. We thank Stefan Pietrzak for scRNA-seq analysis, Dr. Roice Wille for statistical consultation, and the R.S. lab members for critical reading of the manuscript.

## AUTHOR CONTRIBUTIONS

C.K.W. and E.N.N. performed experiments, C.K.W. completed bioinformatic analysis,

C.K.W. and R.S. wrote the manuscript, R.S. acquired funding, A.D. provided reagents and mice,

R.S. and C.K.W. conceived and directed the project, and all authors edited and reviewed the manuscript.

