## Supplemental Figures for "DOT1L interaction partner AF10 controls patterning of H3K79 methylation and RNA polymerase II to maintain cell identity"

### Figure S1

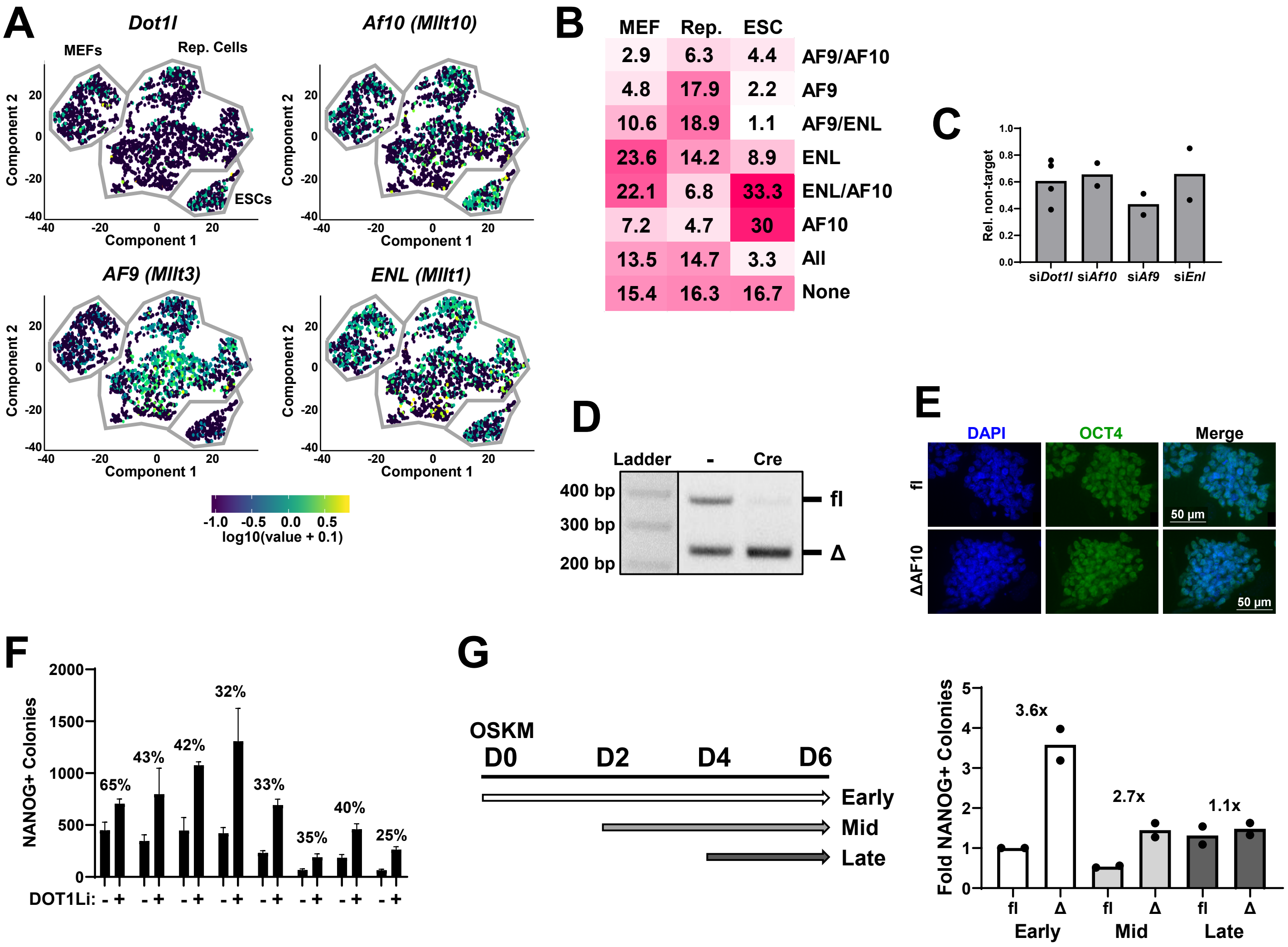


**Figure S1.**

A. scRNA-seq (Tran et al., 2019) t-SNE of MEFs, Reprogramming (Rep.) cells on days 3, 6, 9, and 12, and ESCs. Every cell is represented by a dot and expression of the indicated factor is indicated by color.

B. Percent of DOT1L+ cells from scRNA-seq dataset (Tran et al., 2019) that co-express AF10, AF9, ENL, All, or None of the factors.

C. Relative expression of factors targeted for siRNA depletion on day 3 of reprogramming (Fig 1B). Non-targeting control set to 1. Data are the mean of at least 2 biological replicates, depicted as dots.

D. Addition of adenovirus containing Cre-recombinase deletes the AF10 OMLZ domain from the genome compared to addition of adenovirus containing empty vector control (-), assessed by genotyping PCR. Expected size of the flox (fl) allele = 400 bp, and ΔAF10 (Δ) = 250 bp.

E. Sustained pluripotency was assessed by picking iPSCs at the end of reprogramming, passaging at least 5 times without dox, and staining for pluripotency marker OCT4.

F. Transgene dependent NANOG+ colonies of ΔAF10 with control treatment (-) or DOT1Li (+). Percentages indicate the average colonies in control treated ΔAF10 (DOT1Li treatment set to 100%). Data are the mean (n = 2-3). Each biological replicate depicted individually.

G. Left: AF10 timecourse reprogramming scheme. Cells were treated with control or Cre recombinase adenovirus Early on day 0 (D0), Mid on day 2 (D2), or Late on day 4 (D4) of reprogramming. NANOG+ colonies were analyzed by immunofluorescence on day 6. Right: Reprogramming efficiency of AF10 deletion timecourse. Colonies obtained in fl Early set to 1. Fold calculated relative to the time-matched fl control. Data are the mean (n = 2), biological replicates depicted as dots.

### Figure S2

**A**

| Cluster | # | GO | p-value | Examples |
|---|---|---|---|---|
| **C1** | 10 | N/A | N/A | Osr1, Hoxd12 |
| **C2** | 42 | Negative regulation of cell proliferation | 1.9E-3 | Ndrg2, Sprint1 |
| | | Epithelium development | 2.0E-3 | Epcam, Cdh1 |
| **C3** | 20 | N/A | N/A | Ednrb, Cygb |
| **C4** | 27 | System development | 4.4E-3 | Klf2, Ntn3 |
| **C5** | 10 | N/A | N/A | Aox3, Mmp13 |
| **C6** | 46 | Chemical homeostasis | 3.5E-3 | Esr1, Tmem178 |
| **C7** | 19 | Protein modification process | 3.6E-3 | Nod2, Lck |

**B**

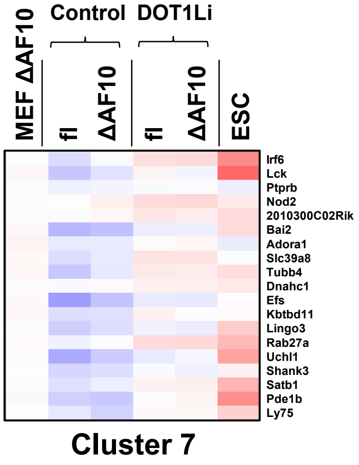


**Figure S2.**

A. Gene Ontology (GO) of each cluster from Fig 2A. GO categories with a p-value less than 0.005 are included.

B. Top: Zoom-in of cluster 7 (Fig 2A).

### Figure S3

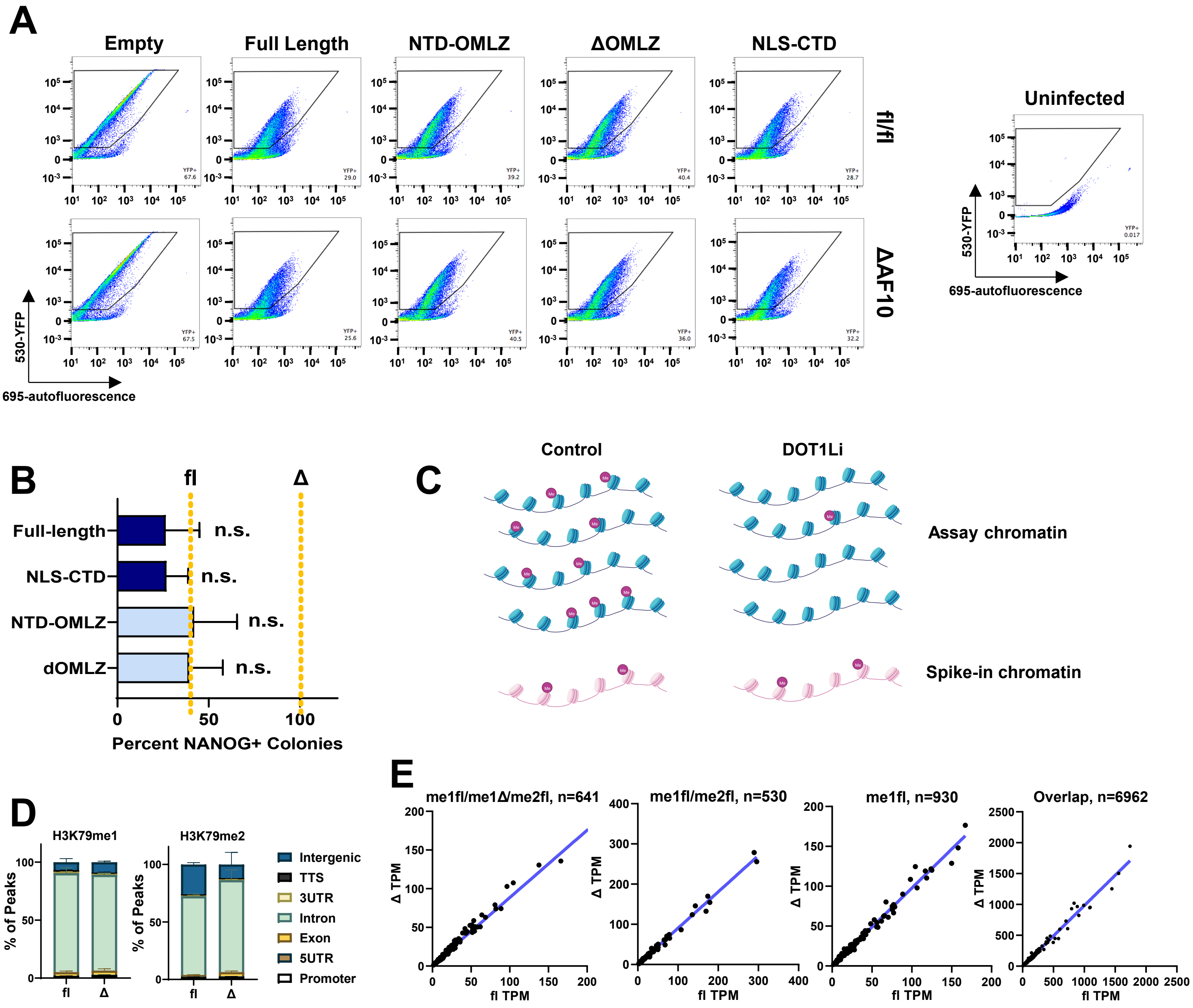


**Figure S3.**

A. Flow cytometry sorting of MEFs infected with AF10 domain mutant-IRES_YFP. Living transduced cells were collected for reprogramming by gating for YFP signal (530 high) compared to uninfected control (Right) with low autofluorescence (695 Low).

B. Reprogramming efficiency of endogenous AF10 wild type (fl/fl) cells expressing the indicated exogenous AF10 domain mutants (Fig 4A). Reprogramming of wild type cells (fl) and ΔAF10 infected with empty vector control indicated by dotted lines. Colonies in ΔAF10 infected with empty vector control set to 100%. Data are the mean + S.D. (n = 4). Not significant (n.s.) P>0.05 by one-way ANOVA.

C. Human spike-in chromatin (293T) was evenly added to all samples before immunoprecipitation. Efficiency of spike-in pull-down was used to scale the assay ChIP-Seq. Created with BioRender.com.

D. Average percent of H3K79me1 and me2 peaks per genomic annotation on day 4 of reprogramming. TTS = transcription termination site and UTR = untranslated region. Data are mean + S.D. (n =2).

E. Expression (TPM) of MEF-specific H3K79me1/2 genes in fl control vs. ΔAF10 on day 4 of reprogramming in the indicated gene lists from Fig. 4E.

### Figure S4

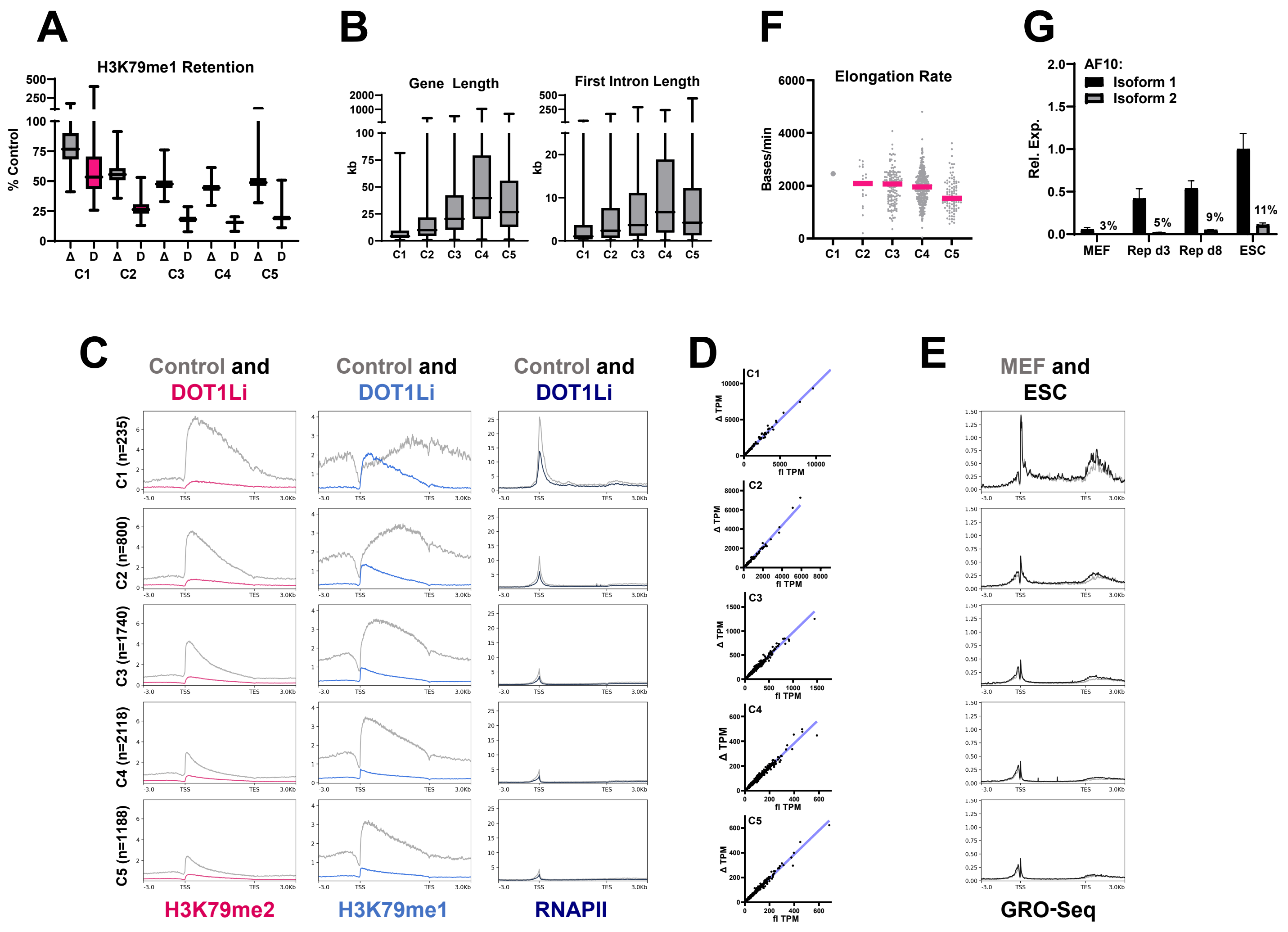


**Figure S4.**

A-F. Clusters of genes with a shared H3K79me1 and me2 peak on day 4 of reprogramming (Fig 5A) were analyzed for:

A. Retention of H3K79me1 enrichment per gene in ΔAF10 (gray - fl set to 100%) and DOT1Li (pink - control set to 100%).

B. Gene length (Right) and first intron length (Left).

C. H3K79me2 (pink), H3K79me1 (blue), and RNAPII (navy) metaplots of DOT1Li and control treated cells on day 4 of reprogramming.

D. Expression (TPM) in fl control vs. ΔAF10 on day 4 of reprogramming.

E. Metaplot of GRO-Seq enrichment in MEFs and ESCs (Min et al., Genes Dev., 2011).

F. Transcription elongation rate in ESCs (Jonkers et al., eLife, 2014).

G. Expression of AF10 (*Mllt10*) isoform 1 (black) and isoform 2 (NM_001252561.1) relative to *Gapdh* in MEFs, reprogramming (Rep), and ESCs. Percentages are the relative amount of isoform 2 (isoform 1 set to 100% per cell type).

### Figure S5

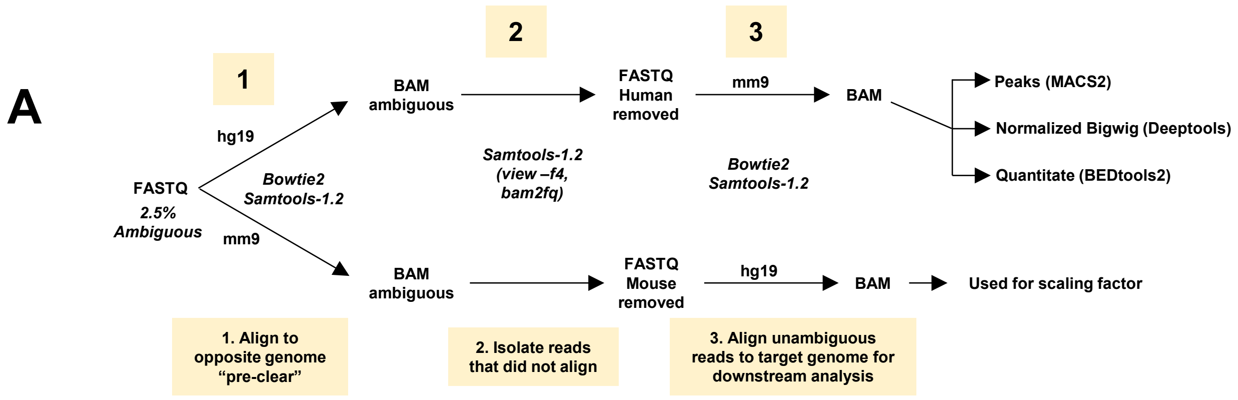


**Figure S5.**

A. Bioinformatic analysis pipeline of quantitative ChIP-Seq. Pre-clearing removed reads that could align to both the mouse and human genome which were about 2.5% of the original FASTQ file.
